# An atypical F-actin capping protein modulates cytoskeleton behaviors crucial to colonization of *Trichomonas vaginalis*

**DOI:** 10.1101/2022.06.09.495492

**Authors:** Kai-Hsuan Wang, Jing-Yang Chang, Fu-An Li, Yen-Ju Chen, Kuan-Yi Wu, Tse-Ling Chu, Jessica Lin, Hong-Ming Hsu

## Abstract

Cytoadherence and consequential migration are crucial for pathogens to establish colonization in the host. In contrast to the nonadherent isolate of *Trichomonas vaginalis*, the adherent one expresses more actin-related machinery proteins with more active flagellate-amoeboid morphogenesis, amoeba migration, and cytoadherence, activities that were abrogated by an actin assembly blocker. By immunoprecipitation coupled with label-free quantitative proteomics, an F-actin capping protein (*Tv*FACPα) was identified from the actin-centric interactome, with an atypically greater binding preference to G-actin than F-actin. *Tv*FACPα partially colocalized with F-actin at the parasite pseudopodia protrusion and formed the protein complexes with α-actin through its c-terminal domain. Meanwhile, *Tv*FACPα overexpression suppresses F-actin polymerization, amoeboid morphogenesis, and cytoadherence in this parasite. Ser^2^ phosphorylation of *Tv*FACPα enriched in the amoeboid stage of adhered trophozoites was reduced by a CKII inhibitor. The site-directed mutagenesis and CKII inhibitor treatment revealed that Ser^2^ phosphorylation acts as a switching signal to alter *Tv*FACPα actin-binding activity and consequent actin cytoskeleton behaviors. Through CKII signaling, *Tv*FACPα also controls the conversion of adherent trophozoite from amoeboid migration to flagellate form with axonemal motility. Together, CKII-dependent Ser^2^ phosphorylation regulates *Tv*FACPα binding actin to fine-tune cytoskeleton dynamics and drive crucial behaviors underlying host colonization of *T. vaginalis*.

## Introduction

*Trichomonas vaginalis* is a pathogenic protist causing trichomoniasis which is one of the most prevalent non-viral sexually transmitted diseases, with approximately 180 million new infections worldwide annually (1).

A successful pathogenic infection includes cytoadherence to establish colonization, followed by migration for population spread. Numerous studies on *Trichomonas vaginalis* have focused on the cytoadherence mechanism in adhesion molecules, like cadherin (2), rhomboid protease (3), legumain protease (4), BAP proteins (5), *Tv*AD1 protein (6), and surface-expressed hydrogenosomal proteins (7, 8, 9, 10). However, the effects of these reputed adhesins in cytoadherence are limited when analyzed by the gain- or loss-of-function assays (2–10). Thus, we postulated that the cytoadherence of *T. vaginalis* might be regulated by pathways other than adhesion molecules. In mammalian adhesion cells, transmembrane integrins link peripheral focal protein complexes underneath the cell membrane for focal adhesion, which is the site that connects the extracellular matrix to transmit traction forces required for cell migration and activates downstream signaling followed by local cytoskeleton reorganization (11, 12, 13). A few studies have used ligand competition or antibody neutralization to demonstrate the involvement of integrin-like molecules in the cytoadherence of *T. vaginalis* (14, 15, 16). Recently, the adherence of clinical *T. vaginalis* isolates to the plastic surface or host cells was shown to be influenced by an actin polymerization blocker (17), implying that the actin cytoskeleton might coordinate cytoadherence in *T. vaginalis*, but the regulatory mechanism was unknown.

Furthermore, flagellate-amoeboid transition immediately after contact with a solid surface or human vagina epithelium cells (*h*VECs), is another striking feature in adherent isolates of *T. vaginalis* (18). Upon morphological transformation, the free-swimming flagellar trophozoite converts to an adherent trophozoite that crawls over a solid surface by pseudopodia-like protrusions referred to as amoeboid migration. A similar flagellate-amoeboid transition was observed in the pathogenic amoeba, *Naegleria fowleri.* This free-living trophozoite builds lamellipodia-like protrusions for phagocytosis and migration driven by actin cytoskeleton machines (19), in which actin expression correlates with its virulence (20).

The actin cytoskeleton is a complex network of actin filaments and actin-associated proteins that shape cell morphology, drive cellular locomotion, and confer cell adhesion (21, 22, 23). The globular actin monomer (G-actin) polymerizes into filamentous actin polymers (F-actin), which are further organized into bundles or branched into three-dimensional networks for complicated cytoskeleton activities. In the polarized F-actin filament, growth initiates from the assembly of the Arp2/3 nucleation complex (24), then G-actin is continuously added at the fast-growing barbed end or dissociated from the pointed end (25). The cellular actin cytoskeleton dynamics are tightly modulated by a variety of accessory effectors for actin polymerization, depolymerization, branching, and reorganization (26). In high eukaryotes, F-actin capping protein (CP) is heterodimerized from α (CPα) and β (CPβ) subunits to form a mushroom-shaped structure capping the fast-growing barbed end of F-actin to block off G-actin access and subsequent polymerization. The C-terminal regions of CPα and CPβ form as two tentacles to bind actin (27, 28, 29). A set of regulatory proteins binds to the barbed end of F-actin to prevent the binding of CP, or several proteins directly bind CP to spatially guide subcellular localization or allosterically alter actin capping activity for instant regulation of cytoskeleton remodeling (30, 31).

Post-translational modifications like phosphorylation and acetylation on the interacting interface within the c-terminal tentacle of CPβ alter the actin-binding dynamics (32). Human CPα forms a protein complex with Casein kinase II-interacting protein (CKIP-1) and Casein kinase II (CKII). CKII phosphorylates the Ser^9^ of CPα coordinating CKIP-1 to inhibit capping activity, but this inhibitory effect seems to be independent of Ser^9^ phosphorylation (33, 34). The capacity of CP binding actin filaments is tightly regulated in a spatial or allosteric manner to fine-tune the actin assembly dynamics in cells.

The mechanisms of actin cytoskeleton regulation in *T. vaginalis* have not been fully elucidated. *Tv*Fimbrin1 protein (*Tv*Fim1) has been identified *in vitro* to accelerate actin assembly and *in vivo* to co-localize with F-actin at the cell membrane periphery in the pseudopod-like structures of *T. vaginalis* upon phagocytosis or migration (35). In this study, a putative F-actin capping protein subunit α (*Tv*FACPα) was identified and characterized from the α-associated protein complexes in *T. vaginalis*.

## Results

### Differential morphogenesis, cytoadherence, and motility of *T. vaginalis*

The differential host-parasite interaction between nonadherent T1 and adherent TH17 isolates was evaluated by the cytoadherence, morphogenesis, and motility. CFSE-labeled trophozoites were co-cultured with the *h*VECs monolayer at MOI of 2:1. Post 60-min infection, ~80% of TH17 but little T1 trophozoites bound to the *h*VECs monolayer (Figure 1A). Most T1 trophozoites maintained an oval-shaped flagellate form, but ~60% of TH17 trophozoites transformed into a flat disk or irregular ameboid form and tightly adhered to the slide surface (Figure 1B). To further observe the dynamics of host-parasite interaction, the trophozoites co-cultured with *h*VECs were monitored by time-lapse imaging (Figure 1C and Videos 1 and 2), showing that nonadherent T1 trophozoites maintained a flagellate form and swam by flagellar locomotion only randomly coming into contact with the *h*VECs. By contrast, adherent TH17 trophozoites rapidly transformed into an amoeboid form within 10 min of contact with the glass slide and crawled toward *h*VECs via pseudopod-like protrusions, referred to as amoeboid migration. In contrast to *T. vaginalis* nonadherent isolate, the adherent isolate displayed more active cytoadherence and amoeboid morphogenesis and migration.

**Figure 1.**
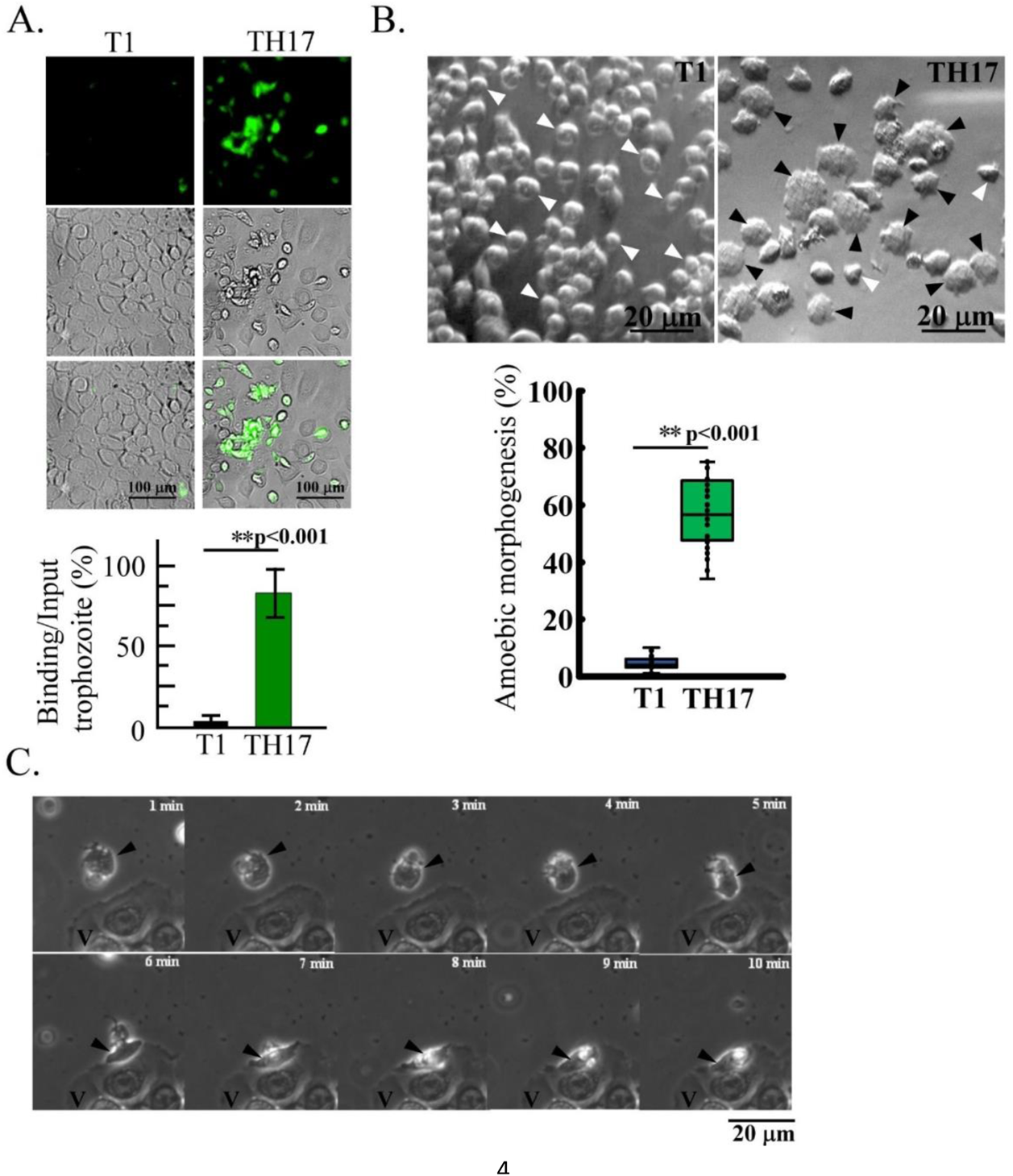
Differential cytoadherence, morphogenesis, migration mode of *T. vaginalis.* A variety of behaviors were observed in the nonadherent (T1) and adherent (TH17) isolates. (A.) The CFSE-preloaded trophozoites were cultured with *h*VECs, then fixed 1 hr post-infection. The cytoadherence capacity was evaluated by the ratio of binding versus input trophozoites as shown in the histogram. Scale bar: 100 μm. (B.) The ratio of T1 or TH17 trophozoites at the amoeboid form was measured in ~600 trophozoites from 12 random microscopic fields as shown in the box and whisker plot. The black and white arrowheads respectively indicate the representative amoeboid and flagellate trophozoites. Scale bar: 20 μm. (C.) TH17 trophozoite (black arrowhead) was co-cultured with *h*VECs (V). The dynamics of ameboid migration and morphogenesis were recorded by time-lapse imaging at one frame per 15 sec over 10 min. Scale bar: 20 μm. All experiments were repeated three times. Data in histogram are presented as mean ± SEM. Statistical significance with p-value for each group of data was analyzed by Student’s t-test as indicated (n=3, *P*< 0.01**, *P*< 0.05*, and ns, no significance).

**(Please see the attached Video 1)**

**Video1 Dynamics of amoeboid morphogenesis and migration in the adherent isolate of *T. vaginalis.*** The trophozoites from TH17 adherent isolate were co-cultured with *h*VECs. The dynamics of trophozoite activities were recorded by time-lapse imaging at the capturing rate of one frame per 30 sec over time as defined.

**(Please see the attached Video 2)**

**Video2 Dynamics of migration in the nonadherent isolate of *T. vaginalis*.** The trophozoites from nonadherent T1 isolate were co-cultured with *h*VECs. The dynamics of trophozoite activities were recorded by time-lapse imaging at the capturing rate of one frame per 30 sec over time as defined.

### Differential expression of actin-related proteins in *T. vaginalis*

Cytoadherence and migration in *T. vaginalis* correlate with actin cytoskeleton (17, 35), therefore, the expression of α-actin and α-actinin, the respective major component and actin bundle linker protein in the cytoskeleton were investigated. The expression of α-actin and α-actinin varied between isolates and was higher in adherent TH17 and T016 isolates compared to nonadherent T1, special in a fresh adherent isolate from a clinical vaginitis patient (Figure 2A). The overexpression of HA-*Tv*actin in a nonadherent isolate did not induce amoeboid morphogenesis or cytoadherence (Figure 2-Figure Supplement 1), suggesting that α-actin might be determinant but insufficient to confer cytoadherence in a nonadherent isolate. Additionally, no detectable α-actin on the adherent parasite surface indicates that α-actin is unlikely to act as an adhesion molecule (Figure 2-Figure Supplement 2).

**Figure 2.**
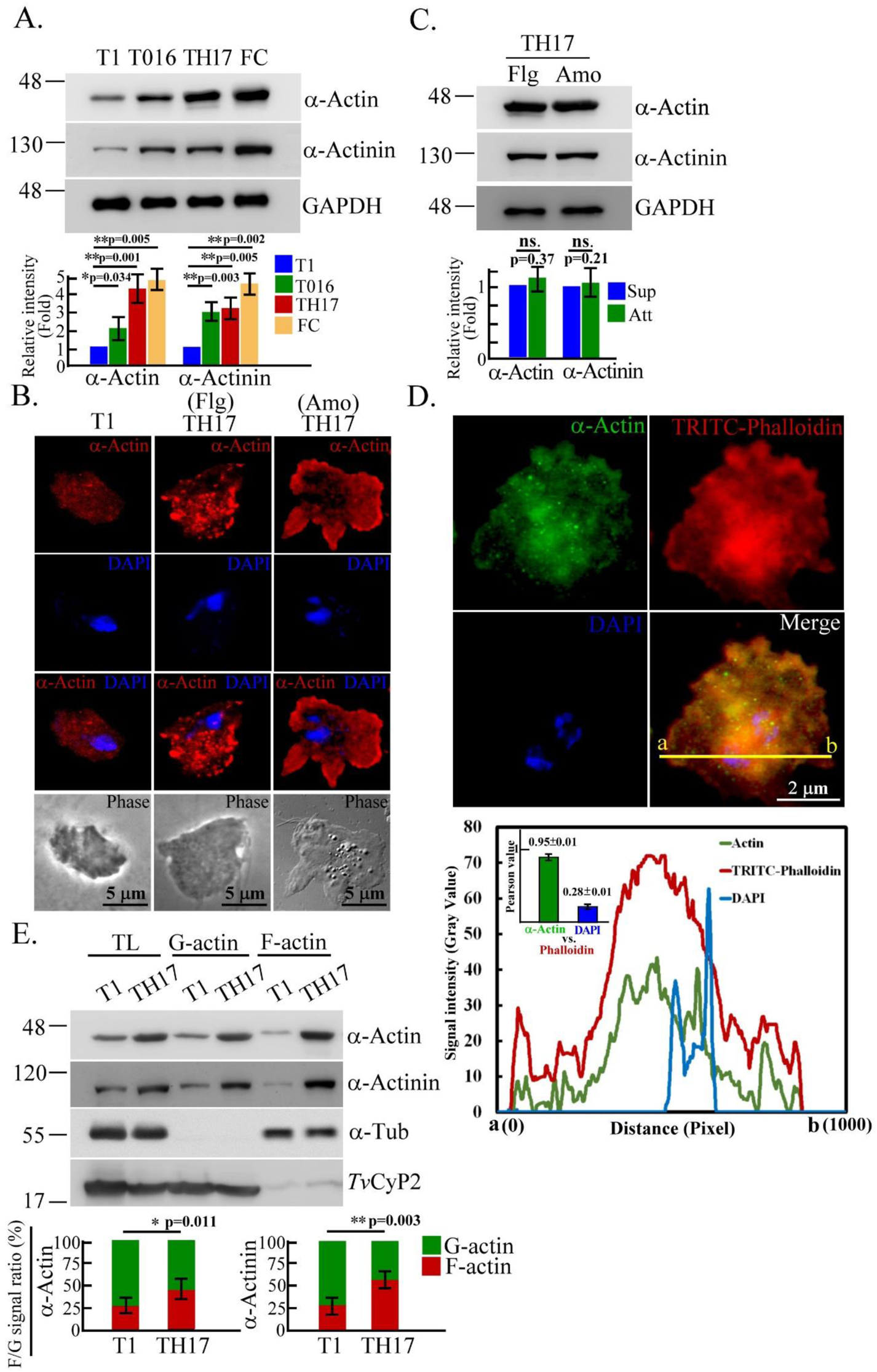
Differential expression of actin-based machinery proteins in *T. vaginalis.* (A.) The total lysates from T1, T016, TH17, and a fresh clinical isolate (FC) were subjected to western blotting. (B., C.) TH17 flagellates (Flg) trophozoites suspended in the medium or amoeboid (Amo) trophozoites adhered to the glass surface were sampled for IFA as shown in (B.) or western blotting as shown in (C.). Scale bar in (B.), 5 μm. (D.) TH17 cultured on a glass slide and fixed for IFA double-staining with anti-α-actin antibody and TRITC-conjugated phalloidin. Signal colocalization was evaluated by a plot profile analysis to show the signal intensity distribution on the yellow line between a and b sites as shown in the diagram. The colocalization of phalloidin with α-actin or DAPI was evaluated by Pearson’s correlation coefficient as shown in the inset histogram. Data are presented as mean ± SEM. Scale bar: 2 μm. (E.) The protein lysates of actin fractionation from T1 and TH17 trophozoites were examined by western blotting. The ratio of indicated protein signal in F-actin and G-actin fractions was analyzed as shown in the histogram. All experiments were repeated three times. Data in histogram are presented as mean ± SEM. Statistical significance with p-value for each group of data was measured by Student’s t-test as indicated (n=3, *P*< 0.01**, *P*< 0.05 **, and ns, no significance).

The immunostaining of α-actin was more intense in TH17 than T1 trophozoites and detected in tiny punctate or short bundles in the cytoplasm of the free-swimming flagellate TH17 but in dense fine networks in the cytoplasm with sporadic clumps underneath the plasma membrane of the amoeboid-adhered TH17. However, the expression of α-actin and α-actinin was similar between the two forms of TH17 trophozoites according to western blotting (Figure 2C). The validated phalloidin binding sites are conserved in α-actin of *T. vaginalis* (Figure 2-Figure Supplement 3) (36). F-actin was double-stained by TRITC-conjugated phalloidin and anti-α-actin antibody (Figure 2D), showing prominent F-actin and α-actin signals concentrated in the juxtanuclear region, referred to as perinuclear actin cap (39), with intense staining underneath the cell membrane of the leading edge in protrusive pseudopods, and less intense staining in the cytoplasm. The signal colocalization of α-actin and phalloidin had a Pearson’s correlation coefficient value of 0.95 (Figure 2D, bottom panel). To evaluate F-actin assembly in cells, G-actin, F-actin, and co-sediments were fractionated, and western blotting analysis revealed an F-actin ratio of ~70% in the adherent isolate and ~30% in the nonadherent one (Figure 2E), similar to α-actinin. It is speculated that F-actin polymerization is more active in the adherent than the nonadherent isolate, and that the actin assembly pattern is also distinct.

### Actin-based morphogenesis, migration, and cytoadherence in *T. vaginalis*

Latrunculin B (LatB) binding sites are conserved in α-actin of *T. vaginalis* (Figure 2-Figure Supplement 3), therefore adherent TH17 trophozoites were treated with LatB to study the role of F-actin in cytoskeleton behavior and cytoadherence. LatB treatment reduced the ratio of F-actin assembly (Figure 3A) and morphogenesis (Figure 3B) in the parasite compared to the DMSO control, as well as decreasing the wound closure rate (Figure 3C) and cytoadherence 60-min post-infection (Figure 3D), showing that F-actin disorder retarded morphogenesis, amoeboid migration, and cytoadherence of this parasite. To rule out the effects from the reputed adhesion molecules (2, 7, 10), the expression of AP65 and PFO (Figure 3E), and their surface distributions (Figure 3F) were analyzed, showing that there was no change in adhesion molecules in the trophozoites with or without LatB treatment. Under IFA permeation condition, hydrogenosomal colocalization of AP65 and PFO proved their surface signal specificities (7, 10). Also, the surface localization of HA-tagged Cadherin-like protein (CLP) was not affected by LatB treatment (Figure 3G). Taken together, actin polymerization is positively associated with the parasite morphological transition, amoeboid migration, and cytoadherence. Also, LatB-inhibited cytoadherence might be independent of adhesion molecules.

**Figure 3.**
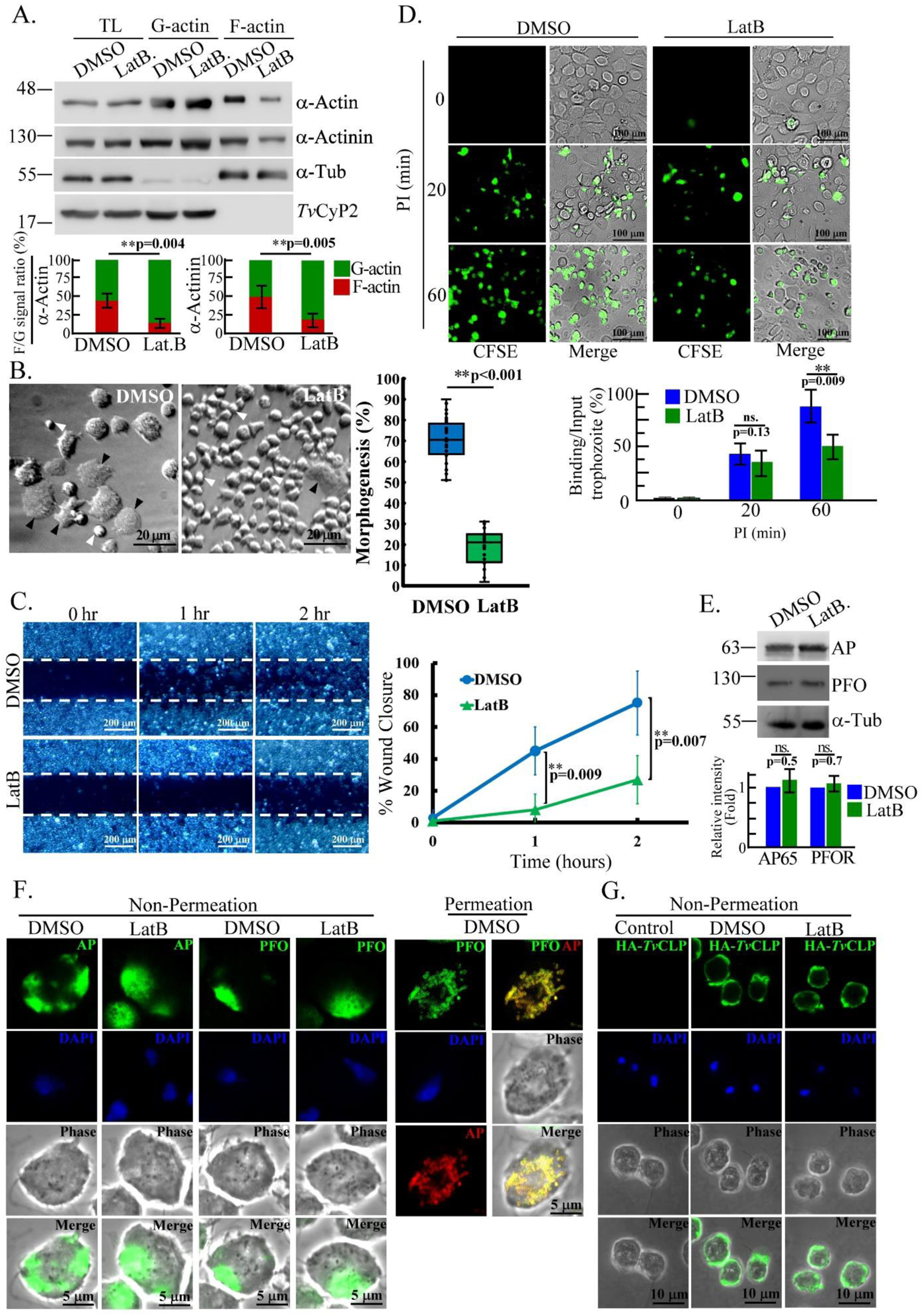
The dysregulation of cytoskeleton-dependent behaviors in *T. vaginalis.* TH17 adherent trophozoites pretreated with DMSO or LatB were sampled for various assays. (A.) Total lysates (TL) or protein lysates of actin fractionation were subjected to western blot. The signal ratio of F-actin versus G-actin (F/G) was measured as shown in the histogram. (B.) Trophozoite morphology was observed by phase-contrast microscopy. The proportion of trophozoites in amoeboid form was measured in 600 trophozoites from 12 random microscopic fields as shown in the box and whisker plot. The black and white arrowheads respectively indicate the representative amoeboid and flagellate forms of trophozoites. Scale bar: 20 μm. (C.) In the wound healing assay, the representative images were captured at 0, 1, and 2 hr. The wound closure rate was measured as the percentage of wound recovery area at indicated time points. The white dashed lines depict the initial wound edge. (D.) In the binding assay, the conditional trophozoites were co-cultured with *h*VECs for the time as indicated. The ratio of trophozoites binding versus input was calculated as shown in the histogram. Scale bar: 100 μm. (E.) Total lysates from conditional trophozoites were subjected to western blotting. (F.) The fixed trophozoites with or without permeation were stained by anti-PFO and anti-AP65 antibodies for IFA. Scale bar: 5 μm. (G.) The non-transgenic control and transgenic trophozoites overexpressing HA-*Tv*CLP were stained by anti-HA antibody for IFA under non-permeation conditions. Scale bar: 10 μm. All experiments were repeated three times. Data in histogram and line chart are presented as mean ± SEM. Statistical significance with p-value for each group of data was analyzed by Student’s t-test as indicated (n=3, *P*< 0.01**, *P*< 0.05 **, and ns, no significance).

### *Tv*FACPα as an α-actin effectors

Since α-actin is not sufficient to promote the cytoadherence in *T. vaginalis* nonadherence isolate, we attempted to identify the regulatory proteins in the α-actin-associated complexes. HA-*Tv*actin was immunoprecipitated from transgenic TH17 trophozoites and subjected to mass spectrometry analysis (Figure 4A), identifying 41 α-actin-associated proteins with an emPAI score above 0.25 or specific in the immunoprecipitant of HA-*Tv*actin (Table 1). These proteins were classified by function into multiple cellular pathways, including cytoskeleton proteins (22%), chaperones (5%), membrane trafficking and transporter (10%), protein binding or modification (7%), DNA/RNA regulation and translation (17%), metabolism enzymes (37%), and uncharacterized proteins (2%) (Figure 4-Figure Supplement 1). The top five abundant protein identified in IP proteome were listed in Figure 4B. Bait HA-*Tv*actin was identified with an emPAI score of ~9.5, an F-actin CP subunit α homolog, referred to as *Tv*FACPα (TVAG_470230), had an emPAI score of ~9.7 (Figure 4B) and 40% identified peptide coverage (Figure 4C), supporting the possibility of a strong protein-protein interaction between *Tv*FACPα and *Tv*actin. The in-silico protein sequence analysis revealed that *Tv*FACPα encodes 267 amino acids with a molecular weight of 29.1 kDa and a PI value of 5.43 and shares 17% identity and 63% similarity with CPα from high eukaryotes (Figure 4D). *Tv*FACPα contains a highly conserved actin-binding domain at C-terminus spanning amino acids from 237 to 261. By a phosphorylation site prediction algorithm (NetPhos 3.1 Generic phosphorylation prediction: Services.healthtech.dtu.dk/service.php?NetPhos-3.1), Ser^2^, Ser^46^, Ser^88^, Ser^106^, and Ser^223^ were predicted as CKII phosphorylation sites. The sequence of ^2^SESE^5^ fits the putative CKII phosphorylation motif (pS/pTDXE) possibly recognized by a phospho-CKII substrate antibody. In the TrichDB database, BLAST analysis identified two CPα homologous proteins (TVAG_470230 and TVAG_212270) with 32% sequence similarity (Figure 4-Figure Supplement 2) but whether they are functionally redundant in this parasite remains to be studied.

**Figure 4.**
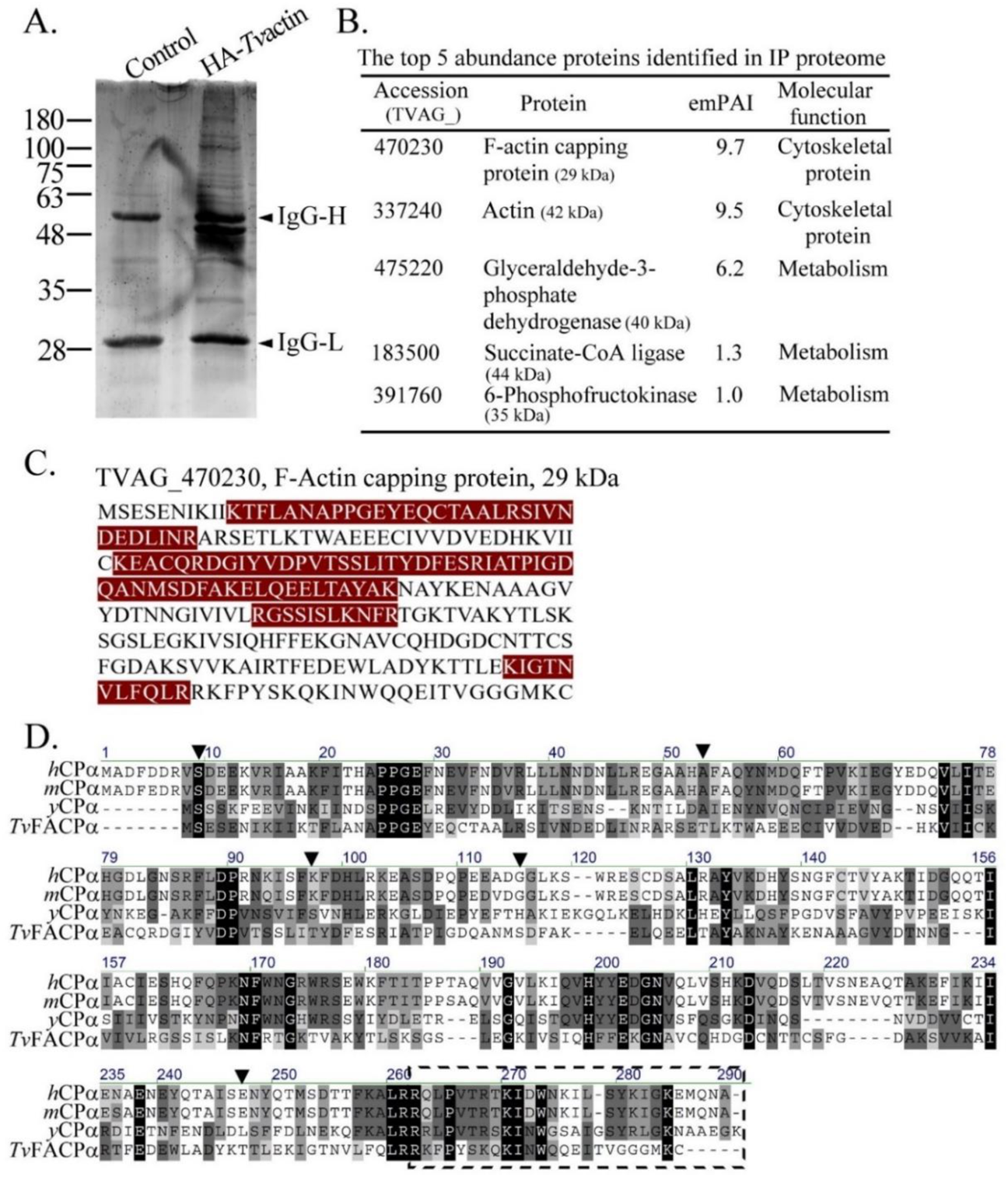
Proteomic identification of actin-binding effectors. (A.) The immunoprecipitants from non-transgenic control or transgenic TH17 trophozoites overexpressing HA-*Tv*actin were separated by SDS-PAGE, followed by SYPRO Ruby staining. (B.) In-gel tryptic digests were processed for a label-free quantitative proteomic analysis. The top five abundant proteins were listed by their emPAI in descending order. The all identified proteins were summarized in Table 1. (C.) *Tv*FACPα-specific peptides identified by LC-MS/MS were labeled in red to show the coverage. (D.) The full-length protein sequence of *Tv*FACPα was aligned with the CPα from human (*h*CPα, P52907), mouse (*m*CPα, P47753), and yeast (*y*CPα, P28495). The conserved amino acid sequences are highlighted. The predicted Casein kinase II phosphorylation sites are indicated by downward arrowheads, and the actin-binding domain is boxed by a black dashed line.

**Table 1.**
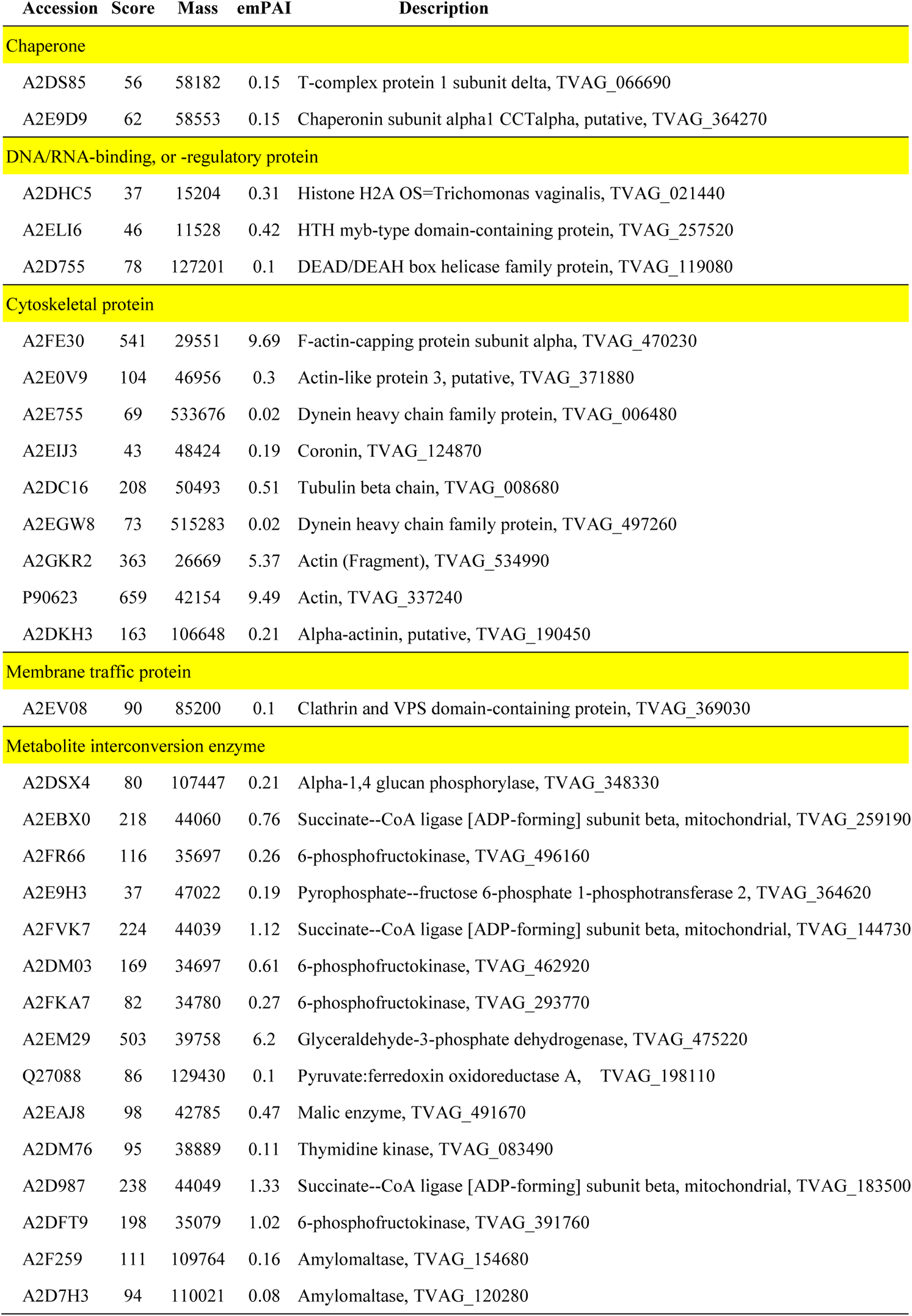
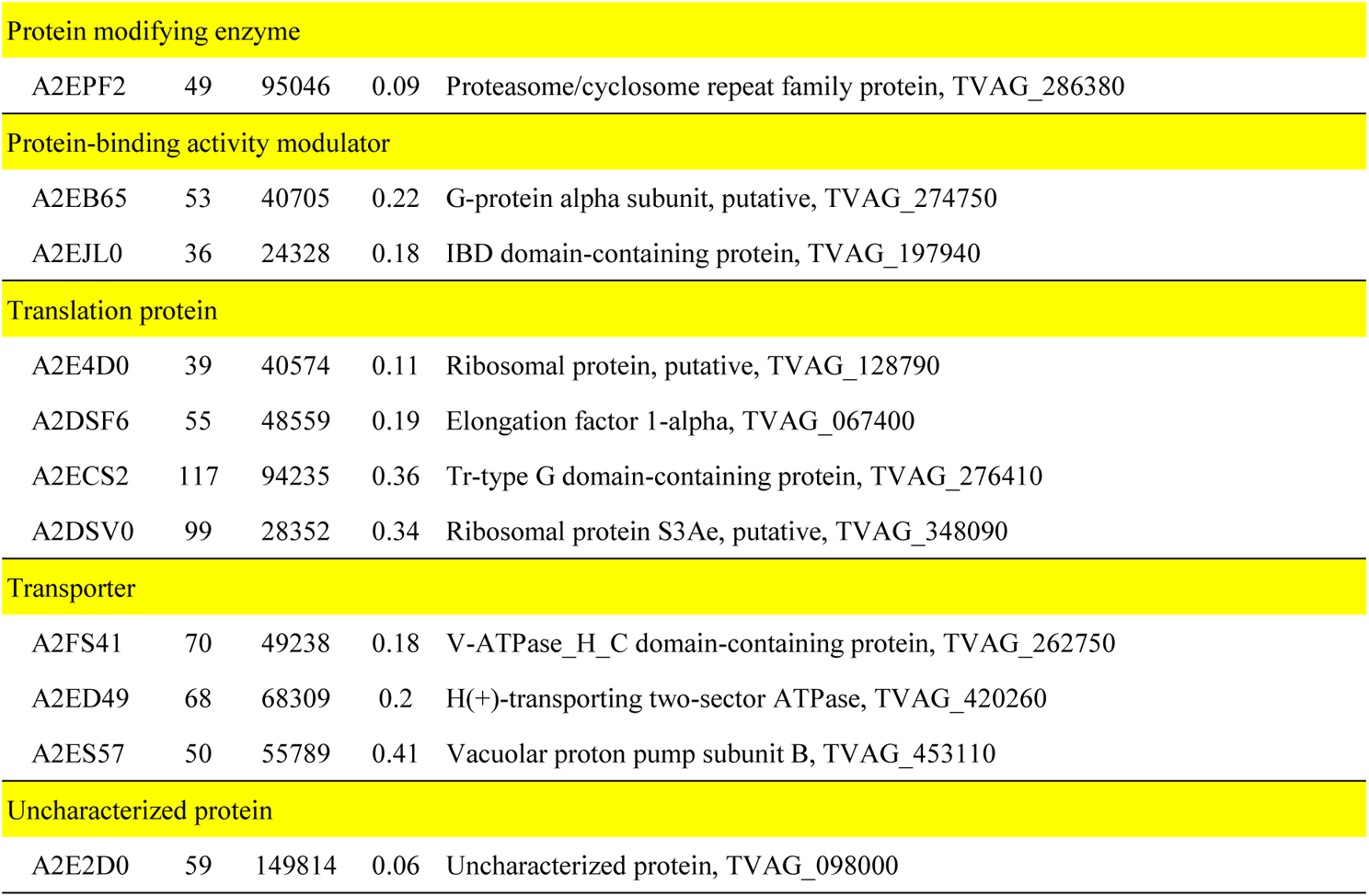
The list of Tvactin-interacted proteins identified by LC-MS/MS. The proteins identified by mass spectrometry with emPAI value above 0.25 or the peptides specific in the immunoprecipitant of HA-*Tv*actin were listed.

| Accession | Score | Mass | emPAI | Description |
| --- | --- | --- | --- | --- |
| **Chaperone** |  |  |  |  |
| A2DS85 | 56 | 58182 | 0.15 | T-complex protein 1 subunit delta, TVAG_066690 |
| A2E9D9 | 62 | 58553 | 0.15 | Chaperonin subunit alpha1 CCTalpha, putative, TVAG_364270 |
| **DNA/RNA-binding, or -regulatory protein** |  |  |  |  |
| A2DHC5 | 37 | 15204 | 0.31 | Histone H2A OS=Trichomonas vaginalis, TVAG_021440 |
| A2ELI6 | 46 | 11528 | 0.42 | HTH myb-type domain-containing protein, TVAG_257520 |
| A2D755 | 78 | 127201 | 0.1 | DEAD/DEAH box helicase family protein, TVAG_119080 |
| **Cytoskeletal protein** |  |  |  |  |
| A2FE30 | 541 | 29551 | 9.69 | F-actin-capping protein subunit alpha, TVAG_470230 |
| A2E0V9 | 104 | 46956 | 0.3 | Actin-like protein 3, putative, TVAG_371880 |
| A2E755 | 69 | 533676 | 0.02 | Dynein heavy chain family protein, TVAG_006480 |
| A2EIJ3 | 43 | 48424 | 0.19 | Coronin, TVAG_124870 |
| A2DC16 | 208 | 50493 | 0.51 | Tubulin beta chain, TVAG_008680 |
| A2EGW8 | 73 | 515283 | 0.02 | Dynein heavy chain family protein, TVAG_497260 |
| A2GKR2 | 363 | 26669 | 5.37 | Actin (Fragment), TVAG_534990 |
| P90623 | 659 | 42154 | 9.49 | Actin, TVAG_337240 |
| A2DKH3 | 163 | 106648 | 0.21 | Alpha-actinin, putative, TVAG_190450 |
| **Membrane traffic protein** |  |  |  |  |
| A2EV08 | 90 | 85200 | 0.1 | Clathrin and VPS domain-containing protein, TVAG_369030 |
| **Metabolite interconversion enzyme** |  |  |  |  |
| A2DSX4 | 80 | 107447 | 0.21 | Alpha-1,4 glucan phosphorylase, TVAG_348330 |
| A2EBX0 | 218 | 44060 | 0.76 | Succinate--CoA ligase [ADP-forming] subunit beta, mitochondrial, TVAG_259190 |
| A2FR66 | 116 | 35697 | 0.26 | 6-phosphofructokinase, TVAG_496160 |
| A2E9H3 | 37 | 47022 | 0.19 | Pyrophosphate--fructose 6-phosphate 1-phosphotransferase 2, TVAG_364620 |
| A2FVK7 | 224 | 44039 | 1.12 | Succinate--CoA ligase [ADP-forming] subunit beta, mitochondrial, TVAG_144730 |
| A2DM03 | 169 | 34697 | 0.61 | 6-phosphofructokinase, TVAG_462920 |
| A2FKA7 | 82 | 34780 | 0.27 | 6-phosphofructokinase, TVAG_293770 |
| A2EM29 | 503 | 39758 | 6.2 | Glyceraldehyde-3-phosphate dehydrogenase, TVAG_475220 |
| Q27088 | 86 | 129430 | 0.1 | Pyruvate:ferredoxin oxidoreductase A, TVAG_198110 |
| A2EAJ8 | 98 | 42785 | 0.47 | Malic enzyme, TVAG_491670 |
| A2DM76 | 95 | 38889 | 0.11 | Thymidine kinase, TVAG_083490 |
| A2D987 | 238 | 44049 | 1.33 | Succinate--CoA ligase [ADP-forming] subunit beta, mitochondrial, TVAG_183500 |
| A2DFT9 | 198 | 35079 | 1.02 | 6-phosphofructokinase, TVAG_391760 |
| A2F259 | 111 | 109764 | 0.16 | Amylomaltase, TVAG_154680 |
| A2D7H3 | 94 | 110021 | 0.08 | Amylomaltase, TVAG_120280 |

| Protein modifying enzyme |  |  |  |  |
| --- | --- | --- | --- | --- |
| A2EPF2 | 49 | 95046 | 0.09 | Proteasome/cyclosome repeat family protein, TVAG_286380 |
| Protein-binding activity modulator |  |  |  |  |
| A2EB65 | 53 | 40705 | 0.22 | G-protein alpha subunit, putative, TVAG_274750 |
| A2EJL0 | 36 | 24328 | 0.18 | IBD domain-containing protein, TVAG_197940 |
| Translation protein |  |  |  |  |
| A2E4D0 | 39 | 40574 | 0.11 | Ribosomal protein, putative, TVAG_128790 |
| A2DSF6 | 55 | 48559 | 0.19 | Elongation factor 1-alpha, TVAG_067400 |
| A2ECS2 | 117 | 94235 | 0.36 | Tr-type G domain-containing protein, TVAG_276410 |
| A2DSV0 | 99 | 28352 | 0.34 | Ribosomal protein S3Ae, putative, TVAG_348090 |
| Transporter |  |  |  |  |
| A2FS41 | 70 | 49238 | 0.18 | V-ATPase_H_C domain-containing protein, TVAG_262750 |
| A2ED49 | 68 | 68309 | 0.2 | H(+)-transporting two-sector ATPase, TVAG_420260 |
| A2ES57 | 50 | 55789 | 0.41 | Vacuolar proton pump subunit B, TVAG_453110 |
| Uncharacterized protein |  |  |  |  |
| A2E2D0 | 59 | 149814 | 0.06 | Uncharacterized protein, TVAG_098000 |

### The non-canonical interaction of *Tv*FACPα to α-actin

Immunoprecipitation was performed to examine whether *Tv*FACPα forms the protein complexes with α-actin in *T. vaginalis*, with two major bands at ~30 and ~32 kDa recognized by an anti-HA antibody in the total lysates. A 42-kDa α-actin and a 110-kDa α-actinin band were co-immunoprecipitated from the trophozoites overexpressing HA-*Tv*FACPα but not the non-transfectant control (Figure 5A). To further confirm the direct interaction of *Tv*FACPα and α-actin, His-*Tv*FACPα, His-△237 mutant, GST and GST-*Tv*actin were purified to homogeneity for the pull-down assay (Figure 5B, left panel). When an equal amount of His-*Tv*FACPα and His-△237 were reacted with GST or GST-*Tv*actin for the pull-down assay, the signal from His-*Tv*FACPα but not His-△237 was pulled down with GST-*Tv*actin, showing that the c-terminal domain is vital for the direct binding of *Tv*FACPα and α-actin (Figure 5B, right panel).

**Figure 5.**
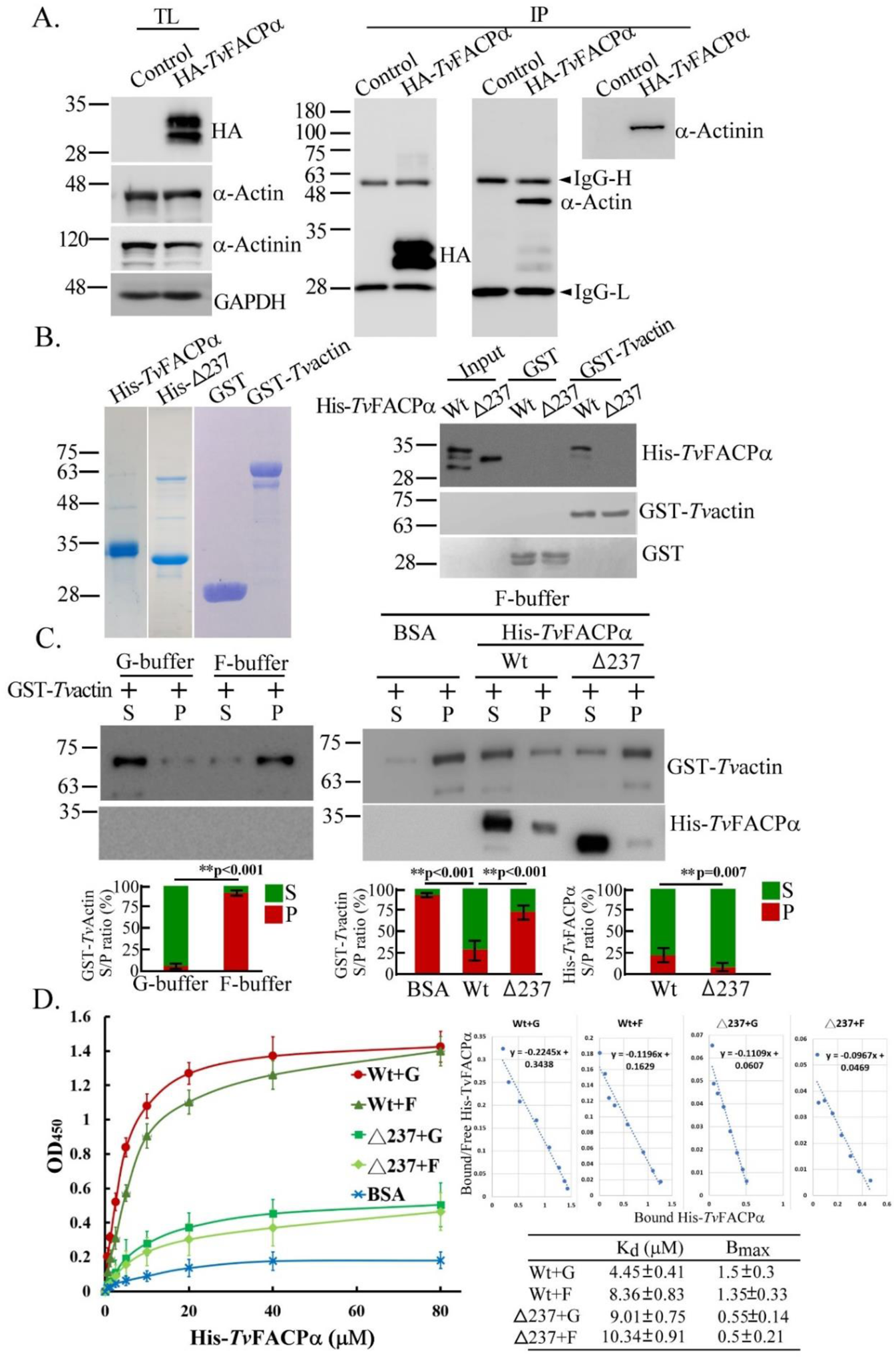
Direct interaction of *Tv*FACPα with G-actin and F-actin. (A.) The total lysate (TL) from non-transgenic control or transgenic TH17 trophozoites expressing HA-*Tv*FACPα was immunoprecipitated by anti-HA antibody (IP), followed by western blotting. GAPDH was detected as the loading control. (B.) The purities of recombinant His-*Tv*FACPα wild type (Wt), His-△237, GST, and GST-*Tv*actin were examined by SDS-PAGE with Coomassie blue staining (left panel). Equimolar GST or GST-*Tv*actin immobilized on glutathione beads was incubated with His-*Tv*FACPα (Wt) or His-△237 mutant for the GST pull-down assay. The pull-down samples were blotted on PVDF membrane for western blotting with anti-6×His antibody or for Ponceau S staining (right panel). 1/10 of the input protein was loaded for positive control. (C.) *In vitro* actin polymerization assay was performed to react G-actin in G- or F-buffer (left panel) or reaction in the presence of His-*Tv*FACPα, His-△237 or BSA control (right panel). After ultracentrifugation, F-actin and its associates were isolated in the pellet (P) from G-actin in the supernatant (S), then examined with western blotting by anti-α-actin or anti-6×His antibody. The signal ratio of indicated protein in the supernatant versus pellet (S/P) was quantified as shown in the histogram. (D.) The equimolar G-actin (G) and F-actin (F) coated on a 96-well microplate were incubated with various concentrations of His-*Tv*FACPα (Wt) or His-△237 (△237) mutant. The saturation binding curve is plotted by the OD_450_ absorbance against various concentrations of His-*Tv*FACPα or His-△237 mutant protein to create Scatchard plots and calculates K_d_ and B_max_ values as summarized in the inset table. The assays were repeated three times. Data are presented as mean ± SEM. Significant difference with p-value for each group of data was statistically analyzed by Student’s t-test as indicated (n=3, *P*<0.01**, *P*<0.05*, and ns, no significance).

The function of *Tv*FACPα in actin assembly was analyzed by an *in vitro* polymerization assay. When over ~95% G-actin polymerized into F-actin in F-buffer in the absence of His-*Tv*FACPα (Figure 5C left panel), F-actin polymerization ratio was only ~25% in the presence of His-*Tv*FACPα, of which 25% of His-*Tv*FACPα co-sedimented with F-actin. By contrast, the polymerization ratio was 75% in the presence of His-△237, and less than 5% of His-△237 could be co-sedimented with F-actin (Figure 5C), indicating that *Tv*FACPα directly interacts with actin molecules to attenuate polymerization. Of note, only 25% of *Tv*FACPα co-sedimented with F-actin but it inhibited over ~70% F-actin formation in the polymerization assay, suggesting that *Tv*FACPα also binds G-actin to inhibit its polymerization.

To determine the kinetics of *Tv*FACPα binding G-actin and F-actin by a solid phase binding immunoassay, two forms of actin were reacted with various concentrations of His-*Tv*FACPα or His-△237 mutant to measure the K_d_ and B_max_ values (Figure 5D). The binding signal increased with increasing concentration of His-*Tv*FACPα or derived mutant, and plateaued in the presence of over 20 μM of His-*Tv*FACPα. The binding curves show that His-*Tv*FACPα binds F-actin with a K_d_ of 8.36 μM and B_max_ of 1.35 and G-actin with a K_d_ of 4.45 μM and B_max_ of 1.5. By contrast, His-△237 binds both F-actin and G-actin with a similar B_max_ of ~0.5, only one-third of His-*Tv*FACPα (Figure 5D inset table). In contrast with the canonical F-actin binding preference for high eukaryotic CPα, the *in vitro* assays demonstrated that *Tv*FACPα bound G-actin with an affinity greater than F-actin to suppress actin polymerization.

### *Tv*FACPα represses F-actin assembly in *T. vaginalis*

By western blotting, the anti-*Tv*FACPα antibody identified a ~30-kDa protein band in the total lysate from TH17 trophozoites (Figure 6A) and colocalized with TRITC-phalloidin with the Pearson’s correlation coefficient value of 0.96, indicating the colocalization of *Tv*FACPα with F-actin in this parasite (Figure 6B). To further study if *Tv*FACPα regulates F-actin polymerization, HA-*Tv*FACPα wild type or actin-binding domain deletion mutant △237 were overexpressed in TH17 trophozoites. By IFA, HA-*Tv*FACPα was detected as network-like structures extending extensively into the cytoplasm and slightly intense immunostaining condensed near the cell membrane (Figure 6C). In the non-transgenic control TH17, α-actin was distributed in the cytoplasm as fine-dense tubular networks. However, cytoplasmic α-actin was observed as numerous stubby rods with punctate signals in HA-*Tv*FACPα-overexpressed TH17, and the pattern in △237 transfectants was similar to that in the non-transgenic TH17 control, indicating that *Tv*FACPα overexpression may alter α-actin organization in this parasite.

**Figure 6.**
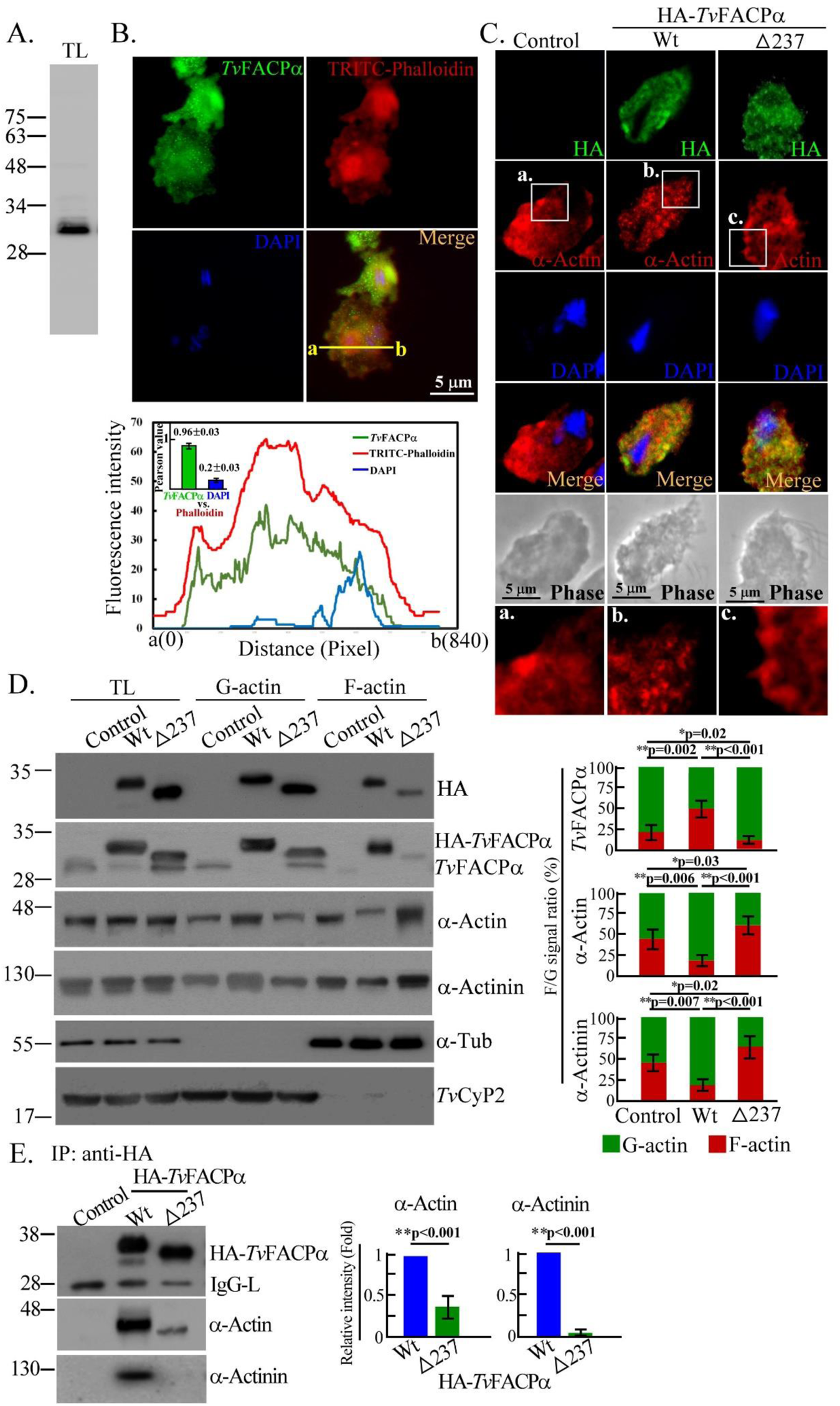
*Tv*FACPα binds actin to block F-actin assembly in *T. vaginalis*. (A.) The total lysate from TH17 trophozoites was subjected to western blotting with an anti-*Tv*FACPα antibody. (B.) TH17 trophozoites cultivated on a glass slide were co-stained with anti-*Tv*FACPα antibody and TRITC-Phalloidin. Signal was assessed by the plot profile analysis to display intensity distribution between sites a to b on the yellow line. The colocalization of phalloidin with α-actin or DAPI was evaluated by Pearson correlation coefficient as shown in the inset histogram. Data are presented as mean ± SEM. Scale bar: 5 μm. (C.) The IFA from the non-transgenic control and transgenic TH17 trophozoites overexpressing HA-*Tv*FACPα or △237 were detected by anti-HA and anti-α-actin. The magnified regions were boxed and images are shown in a-c. Scale bar: 5 μm. (D.) Total lysates and actin fractionations from non-transgenic control and transgenic TH17 trophozoites were examined by western blotting. The ratio of indicated protein signal in F-actin and G-actin fractions (F/G) was analyzed as shown in the histogram. (E.) The total lysates from the trophozoites as shown in (D.) were immunoprecipitated with an anti-HA antibody for western blotting. The relative intensity of indicated protein signal was measured and shown in the histogram. All assays were repeated three times. Data in histogram are presented as mean ± SEM. Statistical significance with p-value for each group of data was analyzed by Student’s t-test as indicated (n=3, *P*< 0.01**, *P*<0.05*, and ns, no significance).

In western blotting, HA-*Tv*FACPα or △237 were overexpressed at a level ~5-fold higher than the endogenous form in the non-transgenic control, and the former inhibited endogenous *Tv*FACPα expression in the transfectant (Figure 6D), suggesting that this parasite may have a feedback mechanism to maintain cellular *Tv*FACPα levels. Western blotting showed that the expression of α-actin or α-actinin did not change between transfectants (Figure 6D). Actin fractionation revealed that ~45% F-actin co-sedimented with ~25% *Tv*FACPα in the non-transgenic TH17 control. In transfectants overexpressing HA-*Tv*FACPα, the F-actin level reduced to ~25% but co-sedimented HA-*Tv*FACPα was ~2-fold higher than the endogenous form of the non-transfectant. In the △237 mutant, the F-actin ratio was slightly higher but co-sedimented △237 was lower than the non-transfectant, therefore, *Tv*FACPα may repress actin polymerization. A similar trend was observed for α-actinin. By immunoprecipitation, co-precipitated α-actin and α-actinin were detected in HA-*Tv*FACPα but much less in the △237 mutant (Figure 6E), indicating that actin-binding activity is essential for *Tv*FACPα to inhibit actin assembly.

### *Tv*FACPα function in actin polymerization is regulated by CKII signaling

Compared to the nonadherent T1 isolate, more *Tv*FACPα and α-actin were detected in adherent TH17 isolates but less *Tv*FACPα co-sedimented with F-actin (Figure 7-Figure Supplement 1). The immunostaining of α-actin was different between the flagellate and amoeboid forms of the adherent isolate (Figure 2B), with equal amounts of *Tv*FACPα, α-actin, and α-actinin detected in the total lysates (Figure 7A). The F-actin ratio in the amoeboid trophozoites was two-fold higher than the flagellate form (Figure 7A), whereas the *Tv*FACPα co-sedimented with F-actin in amoeboid trophozoites was two-fold lower than flagellate form. A similar trend was observed for α-actinin, indicating that adhered-amoeboid *T. vaginalis* displays more active F-actin polymerization and less *Tv*FACPα binding α-actin.

**Figure 7.**
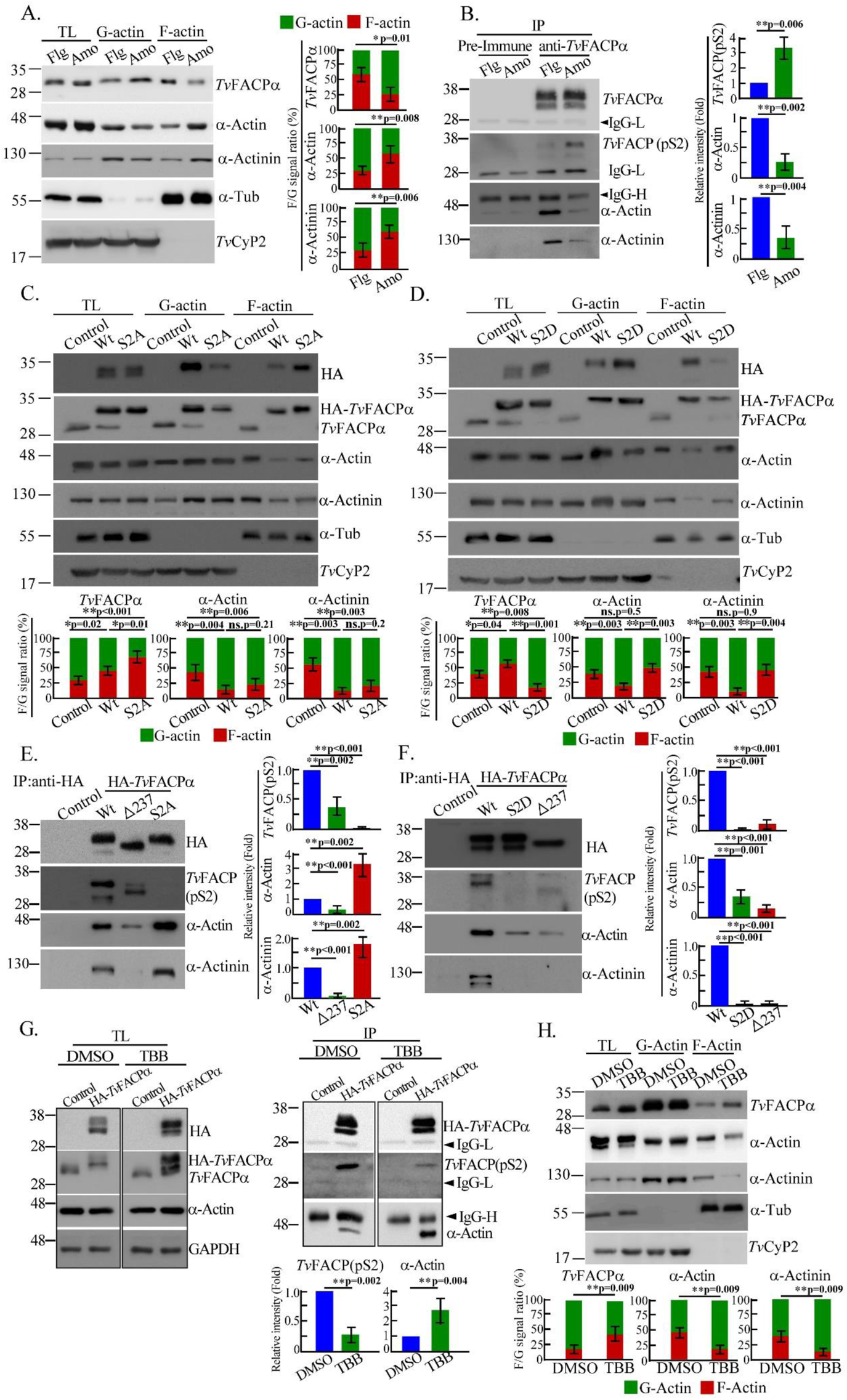
CKII signaling regulates actin-binding of *Tv*FACPα. (A.) Total lysates from TH17 trophozoites in the flagellate (Flg) and amoeboid (Amo) forms were fractionated to determine the ratio of indicated protein signal in F-actin to G-actin (F/G) by western blotting. (B.) The immunoprecipitants from total lysates from (A.) by anti-*Tv*FACPα antibody were examined by western blotting. (C., D.) Total lysates from the non-transgenic control or TH17 trophozoites overexpressing HA-*Tv*FACPα and S2A as shown in (C.), or S2D as shown in (D.), were fractionated for western blotting. The ratio of indicated protein signal from F-actin and G-actin fractions (F/G) was analyzed as shown in the histogram. (E., F.) The total lysates from trophozoites overexpressing HA-*Tv*FACPα and S2A as shown in (E.) or S2D as shown in (F.), were immunoprecipitated by an anti-HA antibody for western blotting. The relative intensities of indicated signals were quantified as shown in the histograms. (G.) The total lysates from the non-transgenic control or HA-*Tv*FACPα-overexpressed TH17 trophozoites with DMSO or TBB treatment were sampled for western blotting (left panel) or immunoprecipitation by anti-HA antibody (right panel). The relative intensities of signals were quantified as shown in the histogram. (H.) TH17 trophozoites treated with DMSO or TBB were fractionated for western blotting. The ratio of indicated protein signal in F-actin and G-actin fractions (F/G) was quantified as shown in the histogram. All assays were repeated three times. Data are presented as mean ± SEM. Statistical significance with p-value for each group of data was measured by Student’s t-test as indicated (n=3, *P*< 0.01**, *P*<0.05**, and ns, no significance).

Regarding the post-translation modifications of *Tv*FACPα, *Tv*FACPα Ser^2^ was previously predicted as a CKII phosphorylation sites (Figure 4D) potentially recognized by a phospho-motif (pS/pTDXE)-specific antibody, referred to as *Tv*FACP(pS2). When *Tv*FACPα was equally immunoprecipitated from the trophozoites, more *Tv*FACP(pS2) but less α-actin and α-actinin were co-pulled down from the amoeboid trophozoites than the flagellate form (Figure 7B). Ser^2^ hyper-phosphorylation enriched in the amoeboid form trophozoites, in which *Tv*FACPα binding α-actin or α-actinin was low. To confirm the role of Ser^2^ phosphorylation in the complex formation of *Tv*FACPα and α-actin, hypo-phosphorylation mimic S2A or hyper-phosphorylation mimic S2D mutant were introduced into TH17 trophozoites for actin fractionation and immunoprecipitation. The overall level of α-actin and α-actinin were similar in the total lysates from *Tv*FACPα, S2A, and S2D transfectants. Compared to the non-transgenic control, both HA-*Tv*FACPα and S2A overexpression repressed F-actin levels in the transfectants, with higher levels of co-sedimented HA-*Tv*FACPα or S2A in the F-actin fraction (Figure 7C). By contrast, a similar level of F-actin was detected in the non-transfectant and S2D mutants but co-sedimented S2D in the F-actin fraction was lower than HA-*Tv*FACPα (Figure 7D). Similar results were obtained for α-actinin. Furthermore, α-actin signals co-immunoprecipitated from the S2A and S2D mutant were three-fold higher and 70% lower respectively than HA-*Tv*FACPα (Figures 7E and 7F), with the low signal intensity of α-actin co-immunoprecipitated with △237 mutant, implying that Ser^2^ phosphorylation is crucial for the actin-binding activity of *Tv*FACPα. Meanwhile, the low intensity of *Tv*FACP(pS2) signal precipitated from △237 mutant implying that the actin-binding domain integrity might be important for Ser^2^ phosphorylation. Ser^2^ phosphorylation is a major signal for the dissociation of *Tv*FACPα and α-actin. The undetectable *Tv*FACP(pS2) signal in the S2A or S2D mutant proves the antibody specificity.

To verify whether Ser^2^ phosphorylation is regulated by CKII signaling, TH17 trophozoites overexpressing HA-*Tv*FACPα were treated with DMSO or TBB for immunoprecipitation and actin fractionation, showing that the overall expression of HA-*Tv*FACPα or α-actin was not influenced by TBB treatment. When an equal amount of HA-*Tv*FACPα was immunoprecipitated from the trophozoites treated with or without TBB, decreasing *Tv*FACP(pS2) but increasing α-actin signals were detected in the co-immunoprecipitants from the parasite treated by TBB (Figure 7G). Furthermore, the overall expression of *Tv*FACPα, α-actin, and α-actinin remained constant in TH17 trophozoites with or without TBB treatment, and when F-actin in TBB-treated parasite was inhibited to one-third of the basal level, the *Tv*FACPα co-sedimented with F-actin was three-fold higher than the DMSO control (Figure 7H). In summary, CKII-dependent Ser^2^ phosphorylation triggers dissociation of *Tv*FACPα and α-actin to evoke actin polymerization.

### *Tv*FACPα in morphogenesis and cytoadherence of *T. vaginalis*

To examine the role of Ser^2^ phosphorylation on cytoskeleton behaviors, the morphogenesis of TH17 trophozoites overexpressing HA-*Tv*FACPα and derived mutants was observed by phase-contrast microscopy. Morphogenesis in the trophozoites overexpressing HA-*Tv*FACPα and S2A was reduced to ~20% compared to ~70% morphogenesis in the non-transgenic control, whereas it was restored to ~70% in the △237 and S2D mutants (Figure 8A). TBB treatment also reduced the morphological transformation of TH17 trophozoites from ~80% in DMSO control cells to ~30% in the TBB-treated trophozoites. Notably, the TBB effect inhibiting morphogenesis was abolished in the S2D transfectant, suggesting that CKII-dependent Ser^2^ phosphorylation in *Tv*FACPα is crucial to the regulation of morphogenesis in *T. vaginalis* (Figure 8B). The differential cytoadherence of various HA-*Tv*FACPα transfectants was monitored over time, showing that the non-transgenic TH17 trophozoites achieved ~100% cytoadherence 60-min post-infection, reducing to ~40% in HA-*Tv*FACPα and S2A transfectants and increasing to ~80% in △237 and S2D transfectants. TBB treatment also significantly reduced the cytoadherence 60-min post-infection and this effect was abrogated in the S2D transfectant (Figure 8D). Notably, the overexpression of HA-*Tv*FACP and related mutants or TBB treatment did not affect the cytoadherence at the initial 20-min infection (Figure 8C). These findings were consistent with our previous observation that LatB only perturbed cytoadherence from the staging 60-min post-infection (Figure 3D). The data strongly supports that CKII-dependent Ser^2^ phosphorylation regulates *Tv*FACPα function in cytoskeleton-mediated morphogenesis and consequential cytoadherence of *T. vaginalis.* The morphogenesis capacity of this parasite tightly correlates its cytoadherence.

**Figure 8.**
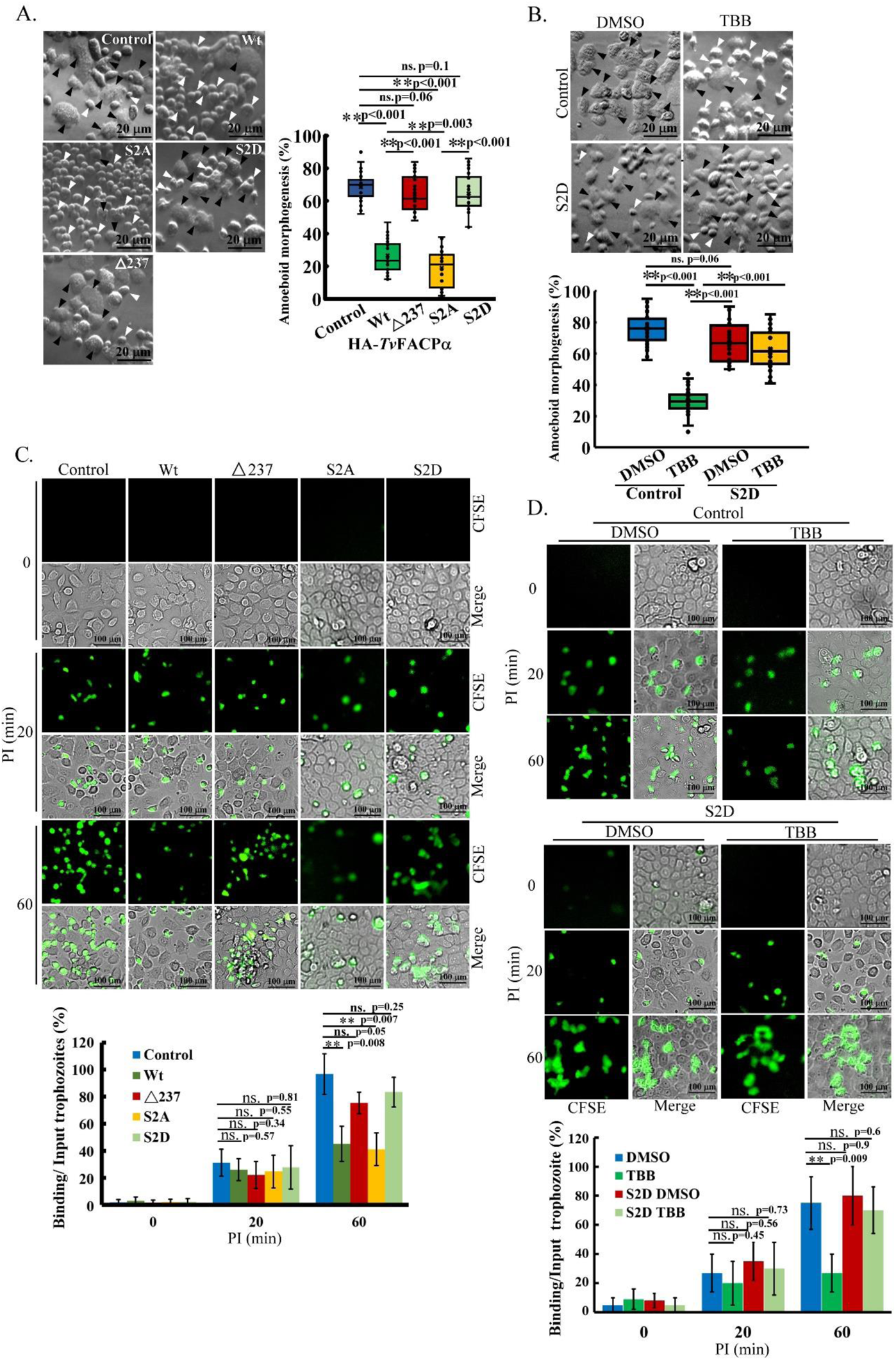
*Tv*FACPα regulates actin-related morphogenesis and cytoadherence in *T. vaginalis*. (A., B.) Non-transgenic control and transgenic TH17 trophozoites overexpressing HA-*Tv*FACPα, △237, S2A, and S2D as shown in (A.), and the non-transgenic TH17 trophozoites and those overexpressing S2D with DMSO and TBB as shown in (B.), were cultured on the glass slide for 1 hr. The cell morphology was recorded by phase-contrast microscopy. The proportion of parasites in amoeboid form was measured in ~600 trophozoites from 12 microscopic fields as shown in the box and whisker plots. The black and white arrowheads respectively indicate the representative amoeboid and flagellate forms of trophozoites. Scale bar: 20 μm. (C., D.) For the cytoadherence binding assay, the CFSE-labeled trophozoites overexpressing HA-*Tv*FACPα and derived mutants as shown in (C.), or the non-transgenic or S2D transgenic TH17 trophozoites pretreated with DMSO and TBB as shown in (D.), were co-cultured with *h*VECs for the indicated timeframes. After removing unbound trophozoites, the ratios of those binding versus input was measured as shown in the histograms. Scale bar: 100 μm. All assays were repeated three times. Data in histogram are presented as mean ± SEM. Statistical significance with p-value for each group of data was analyzed by Student’s t-test as indicated (n=3, *P*< 0.01**, *P*<0.05**, and ns, no significance).

### The function of *Tv*FACPα in amoeboid migration

The conversion of morphology and motility is the dominant features in adherent isolates (Video 1) and retarded by LatB (Figure 3C), so we investigated the role of *Tv*FACPα in amoeboid migration. Since cytoskeletal disorder retarded the morphogenesis and reduced the adherent activity of *T. vaginalis*, the conditional trophozoites had to be sufficiently cultured in the T25 flask until forming a confluent parasite monolayer for the wound heal assay. The wound recovery rate was significantly suppressed in the TH17 trophozoites overexpressing HA-*Tv*FACPα but the rate was similar in the non-transgenic control and △237 mutant, indicating that actin-binding activity is essential for *Tv*FACPα to reduce the amoeboid migration (Figure 9A). Also, the wound recovery rate in the non-transgenic parasite was inhibited by TBB to a similar level to the HA-*Tv*FACPα transfectant. By contrast, the wound closure rate in the S2D mutant was similar to the non-transgenic parasite and not influenced by TBB treatment (Figure 9B), revealing that the S2D mutant counteracted the TBB inhibitory effect on amoeboid migration. This observation indicates that CKII-dependent Ser^2^ phosphorylation might play a key role in *Tv*FACPα-regulated amoeboid migration.

**Figure 9.**
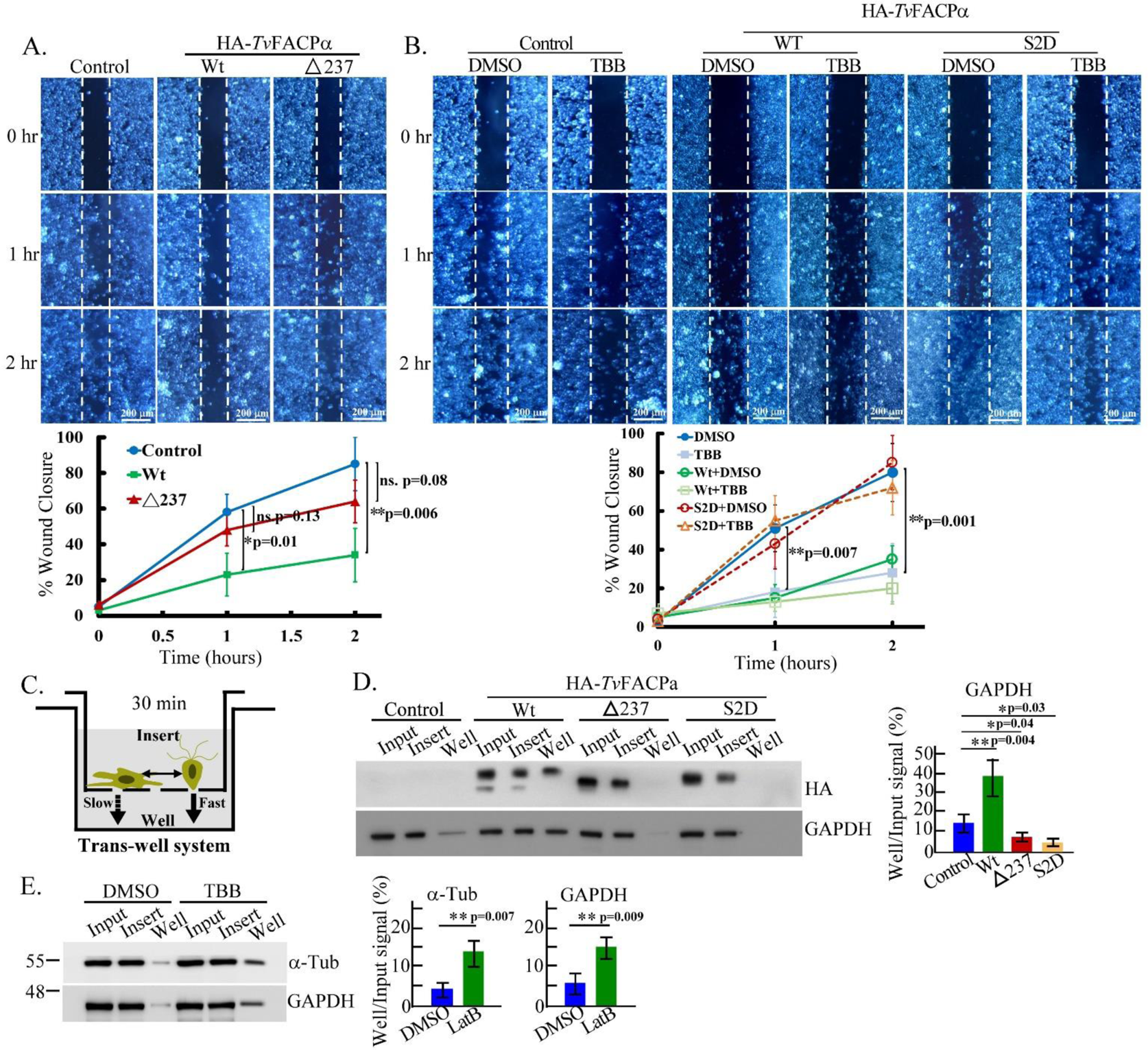
*Tv*FACPα regulates ameboid migration and motility switching of *T. vaginalis*. (A., B.) The migrations of non-transgenic control and TH17 trophozoites overexpressing HA-*Tv*FACPα and △237 in (A.), those overexpressing HA-*Tv*FACPα and S2D with DMSO or TBB treatment in (B.), were evaluated by a scratch wound healing assay. The representative images showing wound closure were captured at 0, 1, and 2 hr, and the closure rate was measured by the percentage of wound recovery area at indicated time points as shown in the line charts. The white dash lines mark the wound scratched boundaries. Scale bar: 200 μm. (C.) A schematic diagram illustrates the working principle of a trans-well system applied to assess migration. Within a short interval, the free trophozoites swim by flagellar locomotion to pass through the boundary membrane faster than crawling by pseudopodia migration. (D., E.) The migrations of TH17 trophozoites overexpressing HA-*Tv*FACPα, △237, and S2D in (D.), TH17 trophozoites treated with DMSO and TBB in (E.), were evaluated by trans-well assay. The relative intensities of signals in the bottom well were evaluated by western blotting and quantified as shown in the histograms. All assays were repeated three times. Data are presented as mean ± SEM. Statistical significance with p-value for each group of data was statistically measured by Student’s t-test as indicated (n=3, *P*< 0.01**, *P*<0.05**, and ns, no significance).

### *Tv*FACPα regulates motility switching in *T. vaginalis*

Next, we tested whether parasite motility is changed with the morphology transition using the trans-well system (Figure 9C). The relative GAPDH signal in the western blotting indicates the relative amount of migratory trophozoites between the bottom wells and top inserts. When GAPDH expression was equal in the input trophozoites, the HA signal was also similar between the transfectants. Focusing GAPDH signal from the bottom well of the 30-min trans-well plate, HA-*Tv*FACPα was higher but △237 and S2D mutants were lower than the non-transgenic control, revealing that more trophozoites with HA-*Tv*FACPα overexpression migrated into the bottom well in a short time (Figure 9D). As observed by microscopy, the trophozoites in the bottom well displayed the morphology at the free-swimming flagellate form (Figure 9-Figure Supplement 1), suggesting that HA-*Tv*FACPα overexpression may retain the parasite in the flagellate form with faster movement driven by motile flagellum. The motility conversion involved actin binding activity regulated by Ser^2^ phosphorylation. TBB inhibited Ser^2^ phosphorylation in *Tv*FACPα (Figure 7G), with the GAPDH signal from the TBB-treated trophozoites in the bottom well higher than in the DMSO-treated trophozoites (Figure 9D). Together, *Tv*FACPα Ser^2^ hypo-phosphorylation retarded amoeboid migration in the adhered trophozoites but expanded the population of free trophozoites that rapidly moved via flagellar locomotion (Figure 9E).

## Discussion

*Tv*FACPα was identified as an actin-binding protein that suppressed actin polymerization via the direct interaction with G-actin monomers and F-actin polymers. Furthermore, CKII-dependent signaling plays a key in the switch from morphology and motility. These cytoskeleton-mediated behaviors are crucial for optimizing the cytoadherence and population spread of this parasite. In the human urogenital tract, the intermittent flushing action of body fluid generates a mechanical barrier to either impair or eliminate the retention of uropathogenic microbes, therefore switching to the opportune motility mode to instantly counteract the environmental challenges or physical defenses would be beneficial for *T. vaginalis* colonization (37).

Unfortunately, the real-time tracking system for fluorescence protein within a living parasite did not work in our assay system, so the overall actin assembly and cytoskeleton activities were evaluated by western blotting and IFA to show the relevance of *Tv*FACPα and actin dynamics in the adherent trophozoites under a steady-state condition.

The DNA sequences of the *tvfacpα* gene from nonadherent and adherent *T. vaginalis* isolates share 100% identity (38), thus the differential cytoskeleton behaviors between isolates are unlikely to be attributed to sequence polymorphism in *Tv*FACPα. Meanwhile, α-actin overexpression dose not promote adherence in the nonadherent isolate, thus cytoskeleton-dependent cytoadherence is unlikely to be determined by one single molecule.

Compared to the nonadherent T1 isolate, more *Tv*FACPα and α-actin were detected in adherent TH17 isolates but less *Tv*FACPα co-sedimented with F-actin (Figure 7-Figure Supplement 1), possibly explaining why the adherent isolate displays more active cytoskeleton behaviors than the nonadherent isolate. Furthermore, the adherent isolate may require a larger *Tv*FACPα reservoir to immediately modulate cytoskeleton dynamics in response to sudden environmental challenges.

The perinuclear actin cap was observed in the trophozoite with dividing nuclei. One of the known functions of the perinuclear actin cap is to govern nuclear location and movement during nuclear division (39), therefore F-actin may function in the nuclear division of this parasite. When there was colocalization of *Tv*FACPα and F-actin at the leading edge of the extending pseudopodia, there was less colocalization observed near the actin cap (Figure 6), suggesting that F-actin bundle assembly in peripheral motile structure is presumably manipulated by *Tv*FACPα, distinct from that in the central juxtanuclear actin cap. Human CKIP-1 protein containing pleckstrin homology domain directs CPα to the cell membrane periphery and bridges the interaction of CPα with CKII kinase to co-regulate cell morphology (33, 34).

*Tv*FACPα Ser^2^ identified as a CKII phosphorylation site is conserved with Ser^9^ on human or yeast CPα (Figure 4D) (33, 34, 40). Human CPα Ser^9^ has been demonstrated to be phosphorylated by CKII kinase but does not directly affect actin assembly (33), indicating that the regulation of human CPα is divergent to *Tv*FACPα in this early evolutionary-branched protozoan. Also, yeast CPα Ser^9^ resides in the stalk domain but not the actin-binding domain, thus Ser^2^ phosphorylation may not directly interfere with *Tv*FACPα actin-binding, instead altering function by an allosteric effect or binding with other CPα interacting partners to co-regulate actin dynamics (40).

Iron was previously found to trigger a protein kinase A-dependent signaling to activate the Myb3 transcription factor sequential phosphorylation and ubiquitination essential to its nuclear translocation (48). However, iron was observed to slightly change *T. vaginalis* morphogenesis long-term cultured in iron-restricted growth medium, so whether iron triggers the CKII pathway to regulate cytoskeleton dynamics in this parasite remains to be elucidated.

When the gain-or loss-of-function assay was employed to study the role of Ser^2^ phosphorylation, F-actin assembly was repressed in the hypo-phosphorylation mimic S2A mutant but restored to near the basal level instead of exceeding it in the hyper-phosphorylation mimic S2D mutant. This implies the existence of additional pathways promoting F-actin assembly under our tested conditions. For example, *Tv*Fim1 protein reveals an opposite function to *Tv*FACPα to accelerate F-actin polymerization that favors phagocytosis and migration in *T. vaginalis* (35). In TrichDB database, BLAST analysis identified two CPα homologous proteins (TVAG_470230 and TVAG_212270) with 32% sequence similarity (Figure 4-Figure Supplement 2) but whether they are functionally redundant in this parasite remains to be studied.

A previous proteomic study reported that surface fibronectin-binding might change actin expression in this parasite. In this report, α-actin expression was constant in the free-swimming flagellate or adhered-amoeboid forms, implying less involvement of fibronectin-binding in the morphogenesis and cytoadherence under our test condition (15).

Mass spectrometry data revealed GAPDH as a major interacting partner of *Tv*actin. In chicken neuron cells, GAPDH acts as a chaperone for α-actin and co-translocates with α-actin to specialized axon sites for polymerization (41). In yeast, GAPDH associates with α-actin and RpB7 subunit of RNA polymerase II to regulate transcription (42, 43). The significance of GAPDH complexed with the actin cytoskeleton in *T. vaginalis* remains to be studied.

The EC_50_ of TBB is varied in different cell types. Numerous CKII alpha subunit (CKIIα) proteins predicted from TrichDB shared less sequence consensus in the TBB binding pocket to high eukaryotic CKIIα (38, 44), possibly explaining why Ser^2^ phosphorylation and downstream cytoskeleton activities were partially inhibited by TBB treatment. Again, the S2D mutation was unable to promote actin polymerization efficiency beyond that of the non-transgenic parasite, suggesting that actin filament growth might be modulated by additional pathways.

The opportunistic amoeba, *Naegleria fowleri,* exists in three life stages: flagellate trophozoite, amoeba trophozoite, and cyst. Multiple environmental factors, like growth temperature, cation level, steroid hormone, or chemical agents, affect the flagellate to amoeba transformation (45, 46, 47). In *T. vaginalis*, other than the contact-dependent effect (14, 18, 35), the factors that trigger the morphological transition are virtually unknown. Overexpression of actin increases the phagocytosis and cytotoxicity of *N. fowleri* (20) but does not affect *T. vaginalis.* (Figure 2-Figure Supplement 1). Although they have the cognate behavior of flagellate-amoeba conversion, their regulation in these two protozoa is distinct. The immediate conversion to motility may allow the parasite to rapidly respond to environmental fluctuations or flushing by humoral fluid flow in the urogenital tract (37).

LatB had little effect on the initial 20-min cytoadherence and surface expression of adhesion molecules, AP, PFO, and cadherin, thus we speculate that the adhesion molecules on the cell surface may play roles in the initial cytoadherence, thereafter actin-based morphogenesis reinforces cytoadherence at the later stage of cytoadherence (49). In our previous study, *Tv*CyP2 was demonstrated to shuttle between intracellular membrane compartments, involving the endoplasmic reticulum, Golgi apparatus, and hydrogenosome before translocation onto the cell membrane (50). The cell surface presentation of adhesion proteins may occur through similar endomembrane trafficking routes.

## Conclusion

In conclusion, *Tv*FACPα directly binds G- or F-actin to block actin filament extension (Figure 10), with Ser^2^ phosphorylation on *Tv*FACPα decreasing actin-binding activity and triggering actin polymerization. In adherent *T. vaginalis* trophozoites, *Tv*FACPα spatially colocalizes with actin molecules at the membrane periphery of motile protrusive pseudopodia, where *Tv*FACPα regulates actin assembly dynamics to control the cytoskeleton behaviors of motility switching, amoeboid migration, or cytoadherence consequent to the morphogenesis. The Ser^2^ phosphorylation status is crucial for *Tv*FACPα function in the regulation of cytoskeleton behaviors. The cytoskeleton-driven activities are also inhibited by a cytoskeleton (LatB) or CKII (TBB) inhibitor. These findings may provide potential therapeutic targets for cytoskeleton aspects to prevent *T. vaginalis* colonization and transmission.

**Figure 10.**
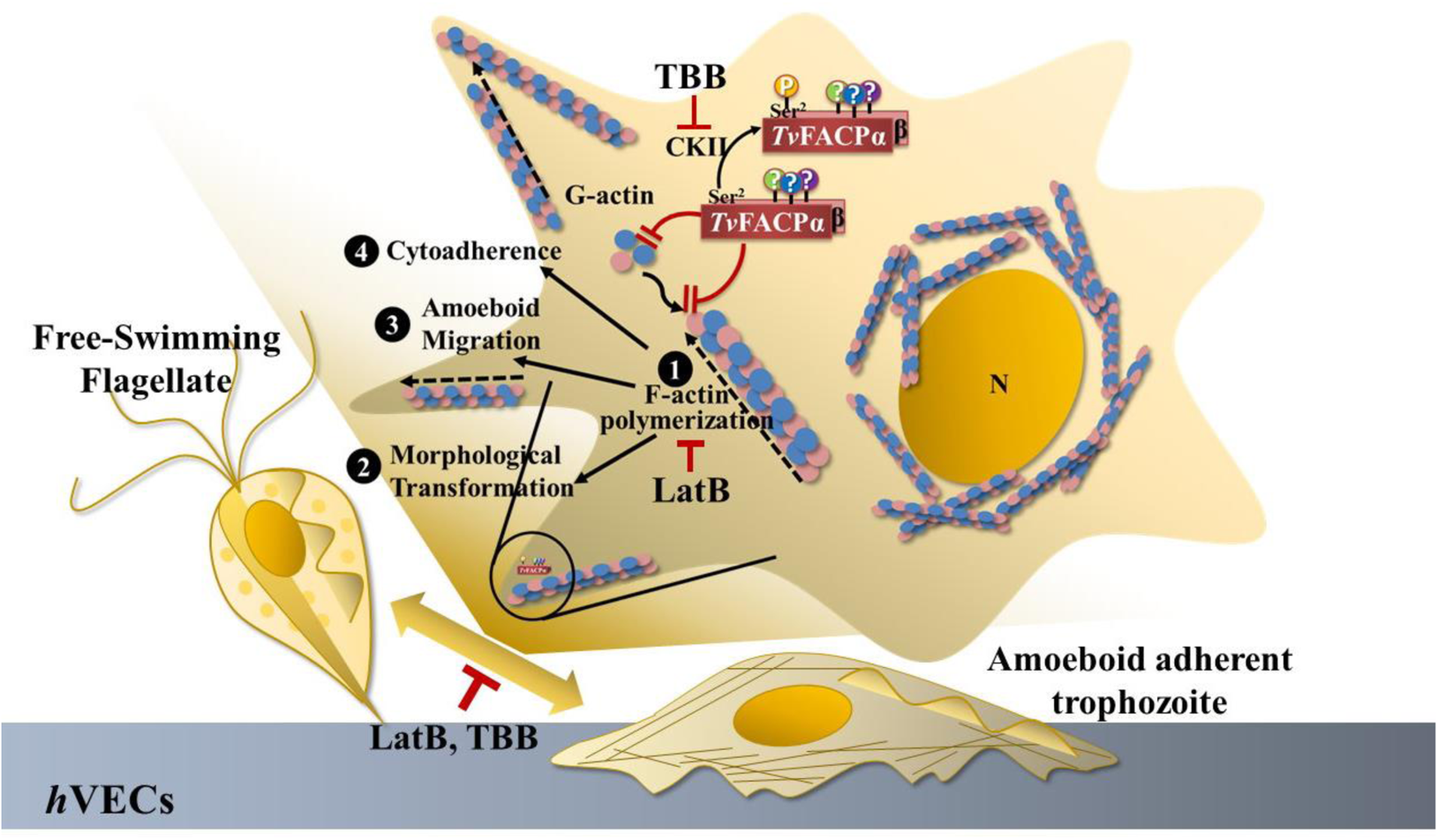
The proposed model for *Tv*FACPα function and regulation. *Tv*FACPα is an actin-binding protein containing a c-terminal actin-binding domain and CKII-dependent Ser^2^ phosphorylation. *Tv*FACPα directly interacts with G-actin and F-actin through the actin-binding domain, and Ser^2^ phosphorylation is the essential signal triggering dissociation of *Tv*FACPα and α-actin. *Tv*FACPα colocalizes with actin at the leading edge of the peripheral motile protrusions inhibiting actin filament polymerization ❶, leading to the diminishment of the flagellate-amoeboid transformation and motility switching ❷, amoeboid migration❸, and the cytoadherence ❹ in this parasite. As expected, the above behaviors were also inhibited by TBB and LatB, supporting the significance of CKII and cytoskeleton activities on parasitism. Tight adherence and immediate migration conversion may be approaches adopted by this parasite to counteract environmental fluctuations or evade the host defense. This novel mechanism of *T. vaginalis* cytoadherence may provide new therapeutic targets for future treatment.

## Materials and Methods

### Cell cultures

*T. vaginalis* trophozoites were cultured in TYI medium at 37°C (48). Two *T. vaginalis* isolates, nonadherent T1 (48) and adherent TH17, were used in this study. T1 with only flagellate trophozoites freely swim in the medium suspension. TH17 displayed vigorous morphogenesis and tightly adhered on glass surface of culture tube. Once the void surface is saturated by adhered trophozoites, the unbound parasite at the flagellate form freely swims in the medium suspension (Figure 1 and Videos 1 and 2). The flagellate trophozoites in the medium suspension and adherent trophozoite on the culture tube surface were collected for analysis as described below. Human vaginal epithelium cells (*h*VECs, VK2/E6E7) were cultivated in Keratinocyte-Serum Free medium (Thermo Fisher Scientific, Massachusetts, USA) at 37°C in 5% CO_2_ as the suggestion by ATCC.

### Lysate preparation from adherent-amoeboid and nonadherent-flagellate trophozoites

Approximately 2×10^7^ trophozoites from TH17 adherent isolate were inoculated into culture tube with 15 ml of medium and incubated at 37°C for 2 hr. The free-trophozoites in suspension were transferred to a new tube and recovered by centrifugation. The cell pellet was lysed in 1 ml lysis buffer (1% Triton X-100, 1× Protease inhibitor cocktail, 1×Phophatase inhibitor cocktail, 100 μg ml^-1^ TLCK, 5 mM EDTA, in TBS). The trophozoites adhering to the glass tube were directly lysed by adding 1 ml lysis buffer and vigorously vortexing for 5 min at 4°C.

### Plasmid construction

The full-length coding sequence of the *tvfacpα* gene (TVAG_470230) was amplified from *T. vaginalis* genomic DNA using the primer pair of *Tv*FACPα-BamHI-5′ and *Tv*FACPα-XhoI-3′. The PCR product was gel-purified, then digested by BamHI/XhoI, and ligated into BamHI/XhoI-predigested Flp-HA-*Tv*CyP2 or pET28a backbone plasmid to obtain Flp-HA-*Tv*FACPα or pET28-His-*Tv*FACPα plasmid. Following a similar procedure, the DNA fragments were amplified from Flp-HA-*Tv*FACP*α* individually using the primer pairs, *Tv*FACPαS2A-5′ and *Tv*FACPα-XhoI-3′ for the S2A mutation, *Tv*FACPαS2D-5′ and *Tv*FACPα-XhoI-3′ for the S2D mutation, and *Tv*FACPα-BamHI-5′ and *Tv*FACPα△237-3′ for the actin-binding domain deletion mutant (△237). The PCR products were gel-purified and subcloned into Flp-HA-*Tv*FACPα or pET28a backbone with BamHI/XhoI sites to generate Flp-HA-*Tv*FACPα(S2A), Flp-HA-*Tv*FACPα(S2D), Flp-HA-*Tv*FACPα(△237), or pET28-His-*Tv*FACPα(△237) plasmid.

To express HA-tagged α-actin in *T. vaginalis* or glutathione S-transferase (GST) fused-α-actin for the GST pull-down or actin polymerization assays, the full-length coding sequence of the *tvactin* gene (TVAG_337240) was amplified from *T. vaginalis* genomic DNA by the primer pair of *Tv*actin-BamHI-5′ and *Tv*actin-XhoI-3′. The gel-purified PCR product was digested with BamHI and XhoI, then ligated into BamHI and XhoI-predigested Flp-HA-*Tv*FACPα or pGST-*Tv*CyP2 plasmid (50) to generate Flp-HA-*Tv*actin, or pGST-*Tv*actin plasmid.

The *Tv*Cadherin expression plasmid was constructed, the coding sequence of the *tvcadherin* gene (TVAG_393390) (2) was amplified from *T. vaginalis* genomic DNA by the primer pair of *Tv*Cadherin-BamHI-5′ and *Tv*Cadherin-XhoI-3′, and subcloned into Flp-HA-*Tv*FACPα backbone vector with *BamH*I and *Xho*I sites to produce Flp-HA-*Tv*Cadherin plasmid.

The primer oligonucleotides used in this study.

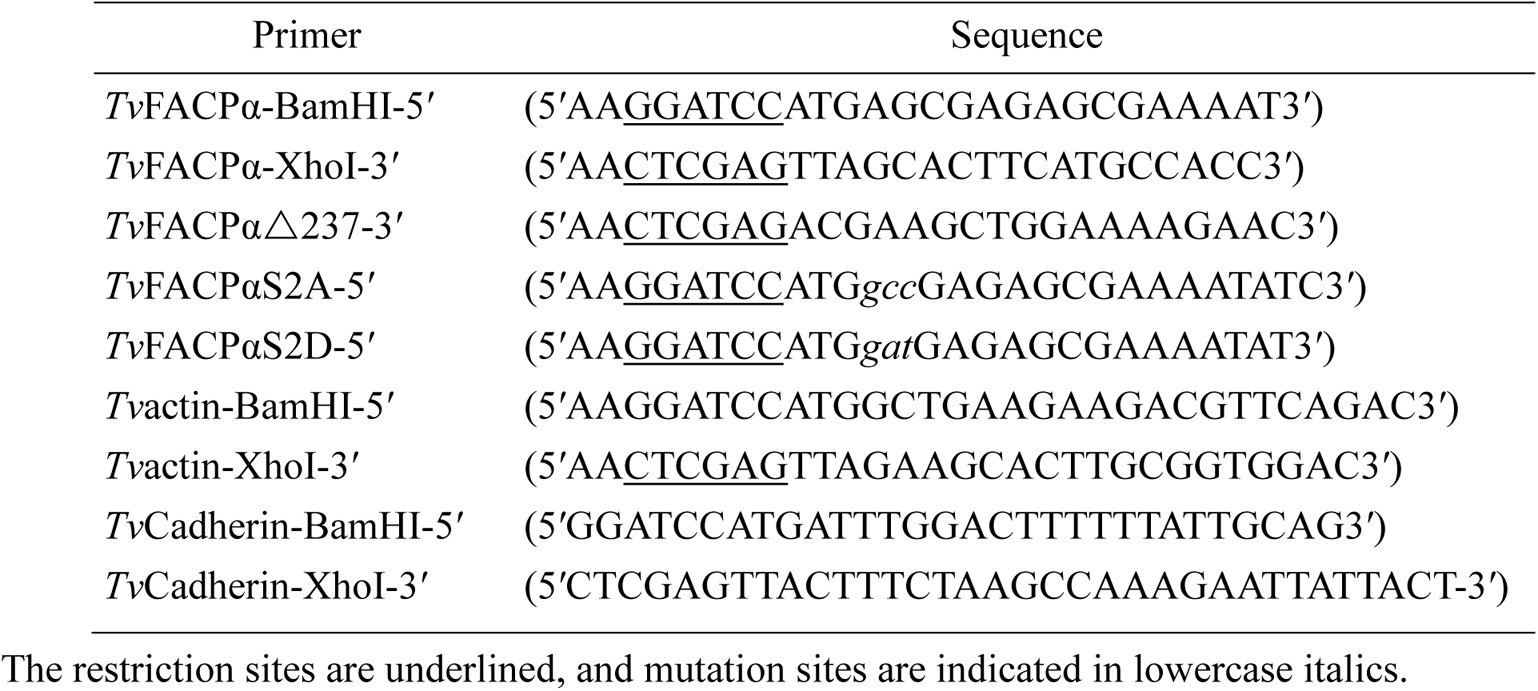

### Cytoadherence binding assay

*h*VECs was cultured in a 24-well plate to an 85% confluent monolayer. Mid-log phase *T. vaginalis* prelabeled with 5 μM of carboxyfluorescein diacetate succinimidyl ester dye (CFSE; CellTrace^TM^, Thermo Fisher Scientific, Massachusetts, USA), were inoculated by a multiplicity of infection (MOI) of 2:1 into *h*VECs culture. At the specific time point, the medium was aspirated and unbound trophozoites were removed by washing two times with PBS for 5 min each. Samples were fixed in 4% formaldehyde for fluorescence microscopy.

### Real-time microscopy

The activity of trophozoites on the confluent *h*VECs monolayer in a glass-bottom culture dish was monitored in real-time by confocal microscopy (LSM-700, Zeiss, Oberkochen, Germany) under a phase-contrast mode with the sampling rate at one frame per 15 sec over time as indicated.

### Inhibitor treatment

1 μM of LatB (Sigma-Aldrich, Massachusetts, USA) or 250 μM of TBB (Sigma-Aldrich, Massachusetts, USA) was added into the *T. vaginalis* culture and incubated at 37°C for 2 hr before analysis.

### Morphology analysis

Trophozoites were cultured on a glass slide in a humid chamber at 37°C for 1 hr and the morphology was observed by phase-contrast microscopy (CKX31, Olympus, Tokyo, Japan). The percentage of flagellate or amoeboid form was measured from 600 trophozoites within 12 random microscopic fields.

### Immunofluorescence assay (IFA)

*T. vaginalis* was fixed with 4% formaldehyde and permeabilized with 0.2% Triton X-100. The samples were then incubated with the primary antibodies: rabbit anti-α-actin (200×, GenScript, New Jersey, USA), mouse anti-α-actin (400×, Abcam Ac-40, Cambridge, UK), mouse anti-HA (200×, Sigma-Aldrich HA-7, Massachusetts, USA), mouse anti-AP65 (7), rabbit anti-PFO (10), rabbit anti-*Tv*FACPα, followed by reaction with FITC or Cy3-conjugated goat anti-mouse or rabbit IgG secondary antibodies (200×, Jackson ImmunoResearch, Pennsylvania, USA). The specimens were air-dried and mounted in medium with DAPI (Vector laboratories, California, USA) for observation by confocal microscopy (LSM-700, Zeiss, Oberkochen, Germany).

### F-actin staining

Trophozoites were fixed with 4% formaldehyde, then permeabilized with 0.2% Triton X-100. The sample was incubated with 20 μg ml^-1^ of TRITC-conjugated Phalloidin (Sigma-Aldrich, Massachusetts, USA) diluted in PBS with 1% BSA in the dark at room temperature for 1 hr. After washing three times with PBS, the glass slide was air-dried and mounted in anti-fade medium (Vector laboratories, California, USA) for fluorescence microscopy (BX-60, Olympus, Tokyo, Japan).

### Signal colocalization evaluation

The fluorescent intensity distributed in the fluorescence assays was measured by plot analysis of ImageJ (Version 1.53q, National Institutes of Health, Maryland, USA). Pearson’s correlation coefficient was calculated to evaluate the signal co-localization, with a value of 1 indicating perfect colocalization, −1 indicating anti-correlation, and 0 representing no correlation.

### Western blotting

The protein samples denatured in 1x SDS sample buffer were separated by sodium dodecyl sulfate-polyacrylamide gel electrophoresis (SDS PAGE) in a 12% gel before blotted to polyvinylidene difluoride (PVDF) membrane by the wet transblot system (Bio-Rad, California, USA). The blocked membrane was incubated with the primary antibodies: mouse anti-HA (2,000×, Sigma-Aldrich HA-7, Massachusetts, USA), mouse anti-α-actin (20,000×, Abcam Ac-40, Cambridge, UK), mouse anti-*Tv*CyP2 (5,000×) (50), mouse anti-α-tubulin (10,000×, Sigma-Aldrich DM-1A, Massachusetts, USA), rabbit anti-*Tv*FACPα (3,000×), mouse anti-6×His (2,000×, Abcam AD1.1.10, Cambridge, UK), rabbit anti-phospho-CKII substrate [(pS/pT)DXE] (1,000×, Cell Signaling Technology, Massachusetts, USA), rabbit anti-PFO (5,000×) (10), mouse anti-AP65 (10,000×) (7), mouse anti-GAPDH (10,000×) (51) and mouse anti-α-actinin (5,000×) (51) at 4°C overnight, followed by HRP-conjugated anti-mouse or rabbit IgG secondary antibodies (5,000×, Jackson ImmunoReaearch, Pennsylvania, USA) at 37°C for 1 hr. The membrane reacted with the enhanced chemiluminescence substrate (ECL, Thermo Fisher Scientific, Massachusetts, USA) were detected and quantified by UVP image system (ChemiDoc-It 815 Imager, VisionWorksLS 8.6 software, Analytik Jena Company, Jena, Germany).

### Immunoprecipitation

Briefly, 6×10^7^ trophozoites were lysed in 1ml of lysis buffer (1% Triton X-100, 1×Protease inhibitor cocktail, 1×Phophatase inhibitor cocktail, 100 μg ml^-1^ TLCK, 5 mM EDTA, in TBS) and centrifuged to remove unbroken cell debris, before the addition of 20 μl of anti-HA antibody-conjugated agarose beads (Sigma-Aldrich, Massachusetts, USA), then incubated on a rotator at 4°C overnight. The beads were recovered by centrifugation and washed three times with 1ml lysis buffer. The precipitates were denatured in 1× SDS sample buffer for western blotting or staining (48, 50).

### Label-free quantitative proteomic analysis

The proteins separated by SDS-PAGE were fixed in methanol for SYPRO Ruby staining (Thermo Fisher Scientific, Massachusetts, USA) and visualization by the Typhoon9410 imaging system (GE healthcare, Illinois, USA). Each gel lane was equally cut into 4 pieces, then sliced into smaller 1-mm^3^ cubes. The gel cubes were desalted by five washes sequentially in 1 ml of 20 mM triethylammonium bicarbonate buffer (TEABC) and 1 ml of 20 mM TEABC with 50% acetonitrile, with the vigorous vortex. The samples were sequentially reduced in 20 mM dithiothreitol (DTT) at 56°C for 1 hr, alkylated in 55 mM iodoacetamide in the dark at room temperature for 30 min and digested with trypsin (Promega, Wisconsin, USA) at 37°C overnight. The tryptic peptides were extracted by vortexing three times sequentially in 20%, 50%, and 100% acetonitrile, then dried in a vacuum concentrator (SpeedVac, Thermo Fisher Scientific, Massachusetts, USA) for LC-MS/MS analysis (48). The protein abundance from the mass spectrometry data was analyzed by a label-free quantitative method by Mascot search, which provides an automated calculation of the Exponentially Modified Protein Abundance Index (emPAI) to estimate the coverage of the identified peptides and abundance for each protein in a dataset. The identified proteins with an emPAI above 0.25 or specific in the co-pull-down sample with their function category are summarized in Table 1. The mass spectrometry proteomics raw data have been deposited to Dryad (https://datadryad.org/stash/share/e30mZQElM-nBNmJOniuiGSBJWBkB7V4-t0XzQ891cX8) or the ProteomeXchange Consortium via the PRIDE (www.ebi.ac.uk/pride/) (52) partner repository with a dataset identifier number of PXD034359.

PRIDE Reviewer access account details:

Username, Password: XpCqEnqW

### In silico analysis of protein sequence and function

The functions of the proteins identified by mass spectrometry were categorized by Protein Analysis Through Evolutionary Relationships (PANTHER) Classification System (www.pantherdb.org/). The *Tv*FACPα protein homologue was searched in TrichDB (trichdb.org/trichdb/app). The multiple protein sequence alignment was analyzed by the Vector NTI AdvanceR 11.5.1 software (Thermo Fisher Scientific, Massachusetts, USA). The protein search was performed by the Basic Local Alignment Search Tool (BLAST, blast.ncbi.nlm.nih.gov/Blast.cgi) or UniProt (www.uniprot.org/).

### Production of recombinant protein

The recombinant protein was produced as previously described (48, 50). The majority of GST-*Tv*actin was expressed in the inclusion bodies of *E. coli* (BL21). For the GST-pull-down assay, the inclusion bodies from 200 ml of *E. coli* culture were dissolved in 1 ml of 8 M urea at 4°C for 20 min to solubilize the proteins. Then, 1 ml of lysate was immediately added to 14 ml PBS and incubated at 4°C for 30 min to refold proteins. After the removal of the insoluble pellets by low-speed centrifugation at 23,000× *g*, soluble GST-*Tv*actin was incubated with glutathione-conjugated sepharose beads as suggested by the supplier (GE healthcare, Illinois, USA) at 4°C for 3 hr and then eluted in GST elution buffer (50 mM Tris-HCl, 10 mM reduced glutathione, pH 8.0). For the solid-phase binding and *in vitro* actin polymerization assays, the bacterial inclusion bodies were solubilized in 8 M urea and directly reconstituted in 14 ml of G-buffer (0.2 mM CaCl_2_, 0.2 mM ATP, 0.5 mM DTT, 5 mM Tris-HCl pH 8.0) at 4°C overnight. The insoluble materials were removed by ultracentrifugation at 100,000× *g* to recover the soluble G-actin in the supernatant (53). Soluble G-actin was further purified by glutathione-conjugated sepharose beads and eluted in G-buffer with 10 mM reduced glutathione.

### GST pull-down assay

GST pull down assay was performed as previously described (50). Briefly, 80 picomoles of GST or GST-*Tv*actin immobilized on 20 μl of glutathione conjugated-sepharose beads 4B (GE healthcare, Illinois, USA) was incubated with 80 picomoles of His-*Tv*FACPα or derived mutants in 1 ml GST binding buffer (PBS contains 0.2% Triton X-100 and 1 mM EDTA) at 4°C rotation overnight The GST beads were washed three times by 1 ml of GST binding buffer then denatured in 1× SDS sample buffer for further analysis.

### *Tv*FACPα antiserum production

The recombinant His-*Tv*FACPα full-length protein was produced and purified by a standard protocol as suggested by the supplier (QIAGEN, Hilden, Germany) (48, 50). Using the purified His-*Tv*FACPα protein to immunize rabbits for antiserum production is a customized service provided by the manufacturer (Genetex, California, USA). The antibody specificity of anti-*Tv*FACPα was tested by western blotting as shown in Figure 6A.

### *In vitro* actin polymerization and co-sedimentation assay

Insoluble GST-*Tv*actin denatured in 0.5 ml of 8M urea was reconstituted in 7.5 ml of G-buffer (0.2 mM CaCl_2_, 0.2 mM ATP, 0.5 mM DTT, 5 mM Tris-HCl, pH 8.0) at 4°C overnight to ensure that the thorough GST-*Tv*actin depolymerizes into the G-actin form. Then, 80 picomoles of G-actin in 1 ml of G-buffer were added 1/10 volume of 10× F-buffer (500 mM KCl, 20 mM MgCl_2_, and 10 mM ATP, 100 mM Tris, pH 7.5) at 4°C for 1 hr to trigger actin polymerization. F-actin was recovered from the pellet by 100,000× *g* ultracentrifugation, whereas soluble G-actin in the supernatant (54). The ratio of F-actin versus G-actin was evaluated with Coomassie blue staining or Western Blot detection. Alternatively, F-actin and co-sediments were recovered from the pellet by 100,000× *g* ultracentrifugation of the *in vitro* actin polymerization assay in the presence of 80 picomoles of His-*Tv*FACPα wild type or His-△237 at 4°C for 1 hr.

### ELISA-based solid-phase binding assay

The solid-phase binding assay was performed as described previously, with a few modifications (54). Briefly, 100 μl of 2.5 μM G-actin in G-buffer or F-actin in F-buffer was added to a 96-well microplate and incubated at 4°C with gentle shaking for 8 hr. After three washes with PBST (0.05% Tween 20 in PBS), the samples were blocked in PBST with 5% non-fat milk at 37°C for 2 hr before 100 μl aliquots of different concentrations of His-*Tv*FACPα (0, 2.5, 5,10, 20, 40, and 80 μM) were added to the wells and incubated at 4°C with gentle agitation overnight for protein-protein interaction. Unbound protein was removed by three washes with PBST and the plate was incubated with mouse anti-6×His primary antibody (10,000×, in PBST containing 5% non-fat milk) at room temperature for 2 hr, followed by three washes with PBST. The wells were incubated with HRP-conjugated goat anti-mouse IgG secondary antibody (5000× in PBST containing 5% non-fat milk, Jackson ImmunoReaearch, Pennsylvania, USA) at room temperature for 2 hr. The wells were washed before the addition of 100 μl/well of 3, 3’, 5, 5’-tetramethylbenzidine (TMB, Sigma-Aldrich, Massachusetts, USA) substrate at room temperature for 5 min. The colorimetric reaction was stopped by the addition of 100 μl/well 1N HCl and the absorbance was detected by spectrophotometry at OD_450_ (Molecular Device, California, USA). The absorbances at OD_450_ were plotted against the concentrations of His-*Tv*FACPα to generate Scatchard plots and calculate K_d_ and B_max_ values (54).

### Actin biochemical fractionation

G- and F-actin were fractionated and enriched using a commercial *in vivo* assay biochem kit (Cytoskeleton Inc, Colorado, USA), according to the manufacturer’s instructions with minor modifications. Briefly, around 3× 10^7^ trophozoites were incubated in cell lysis buffer (Cytoskeleton Inc, Colorado, USA) with vigorous agitation at 4°C for 30 min and homogenized by a 23-gauge needle on a 5-ml syringe. The total lysate was centrifuged at 1,000× *g* to remove the unbroken cell debris, followed by ultracentrifugation at 100,000× *g* for 1 hr to separate the insoluble F-actin and associated proteins in the pellet from soluble G-actin in the supernatant. In western blotting, α-tubulin and *Tv*CyP2 were respectively detected as purity markers for F-actin and G-actin fractions.

### Cell migration assay

For the wound healing assay, adherent *T. vaginalis* trophozoites were cultured to a confluent monolayer in a T25 flask. A scratch (200-μm to 1-mm wide) was generated by scraping the trophozoite monolayer with a P200 tip. After removal of cell debris by washing once with the growth medium, the culture flask was incubated at 37°C and images were captured in a defined area at an interval of 30 min over 2 hr. The wound closure area in each image was measured by ImageJ software (Version 1.53q, National Institutes of Health, Maryland, USA). For the trans-well migration assay, ~1×10^7^ trophozoites suspended in 2 ml of TYI medium were inoculated into the top insert divided by a polyester membrane with 3-μm pores (4.6 cm^2^, JET Biofil, Guangzhou, China). The top insert was placed in a 6-well culture plate containing 2 ml of TYI medium and cultured at 37 °C for 30 min. The trophozoites in the top insert and bottom well were collected for microscopic observation and western blotting.

### Statistical analysis

Statistical significance of data collected from control and conditional samples was analyzed by Microsoft Office Excel 2019 software with Student’s t-test. P< 0.05 is considered as significant difference.

## Acknowledgement

We are grateful to Dr. Jung-Hsiang Tai (Institute of Biomedical Sciences, Academia Sinica, Taiwan) for *T. vaginalis* T1 isolate, Dr. John Alderete (Washington State University, USA) for the anti-α-actinin, anti-GAPDH, and anti-AP65 antibodies, and Dr. Rossana Arroyo (CINVESTA, Mexico City, Mexico) for the anti-PFO antibody. Also, we are grateful to the Proteomics Core Facility (Institute of Biomedical Sciences, Academia Sinica, Taiwan) for the LC-MS/MS analysis. This work was supported by grants from the Ministry of Science and Technology of Taiwan (110-2320-B-002-048-and 110-2320-B-002-076-).

## Author contributions

Kai-Hsuan Wang: Investigation, Validation, and Methodology.

Jing-Yang Chang: Investigation, Validation, and Methodology.

Fu-An Li: Investigation, Validation, and Methodology.

Yen-Ju Chen: Investigation, Validation, and Methodology.

Kuan-Yi Wu: Investigation, Validation, and Methodology.

Tse-Ling Chu: Investigation and Validation

Jessica Lin: Investigation and Validation

Hong-Ming Hsu: Investigation, Validation, Project Administration, Supervision, Funding Acquisition, Conceptualization, Writing-Original Draft Preparation, and Writing-Review & Editing.

## Statement of conflict of interest

The authors declare that they have no competing interests in this manuscript.

## Data availability Section

All data generated or analyzed during this study are included in the manuscript and supplementary data; Source Data files have been provided for Table 1, the statistical analysis of quantification, and raw gel or blot images generated in this study. The proteomics raw data have been deposited to Dryad (https://datadryad.org/stash/share/e30mZQElM-nBNmJOniuiGSBJWBkB7V4-t0XzQ891cX8) and PRIDE (www.ebi.ac.uk/pride/) with a dataset identifier number PXD034359. (PRIDE Reviewer access account details: Username, Password: XpCqEnqW).

## Supplement data and legends

**Figure 2-Figure Supplement 1.**
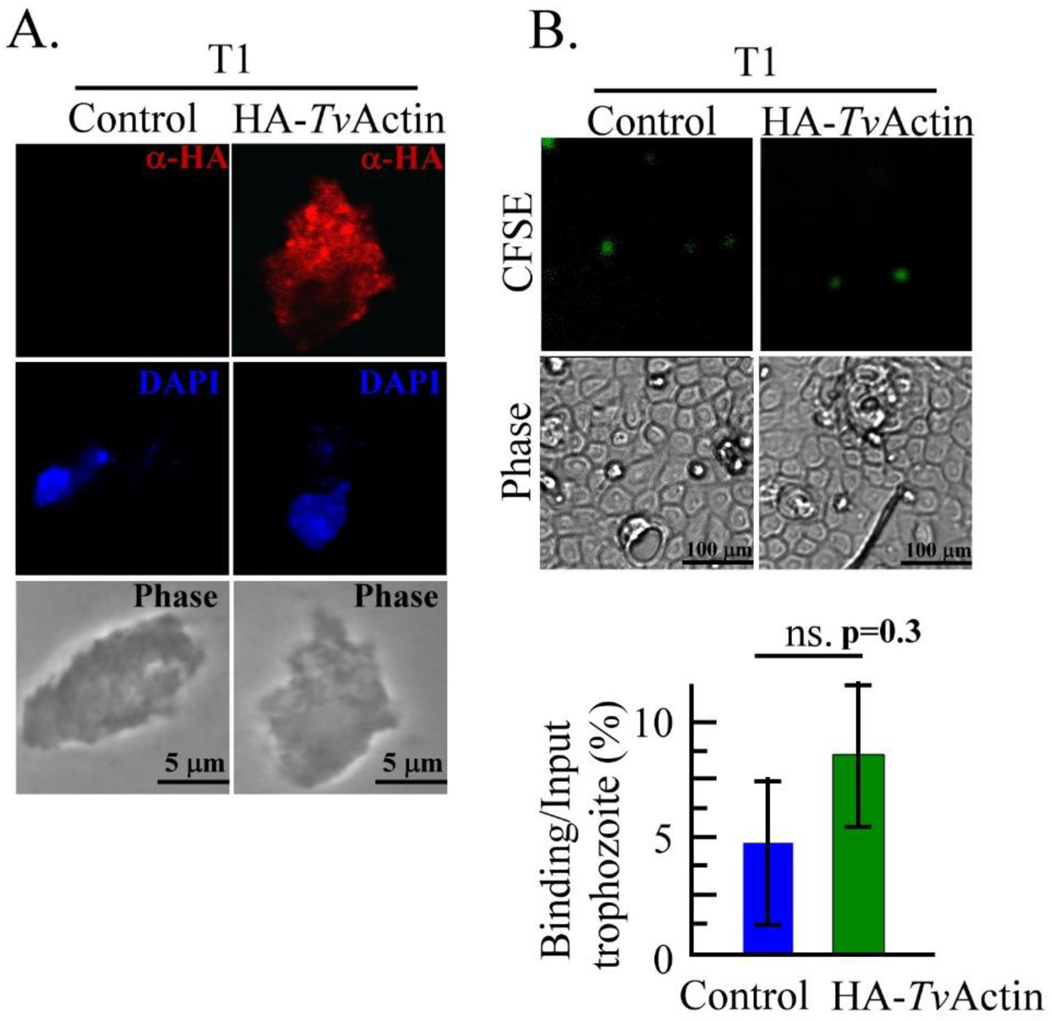
Overexpression of HA-*Tv*actin in nonadherent T1 isolate. (A.) The non-transgenic and HA-*Tv*actin transgenic T1 trophozoites were fixed for IFA by an anti-HA antibody and then incubated with a Cy3-conjugated secondary antibody. The nuclei were stained by DAPI and the morphology was observed by phase-contrast microscopy. The scale bars represent 5 μm. (B.) The non-transgenic and HA-*Tv*actin transgenic T1 trophozoites prelabeled by CFSE were co-cultured with *h*VECs at MOI of 2:1 for 1 hr in the cytoadherence binding assay. The scale bars represent 100 μm. The assays were repeated three times. Data in the histogram are presented as mean ± SEM. Significance with p-value is statistically analyzed by Student’s t-test as indicated. (n=3, P<0.01, P<0.05, and ns, no significance).

**Figure 2-Figure Supplement 2.**
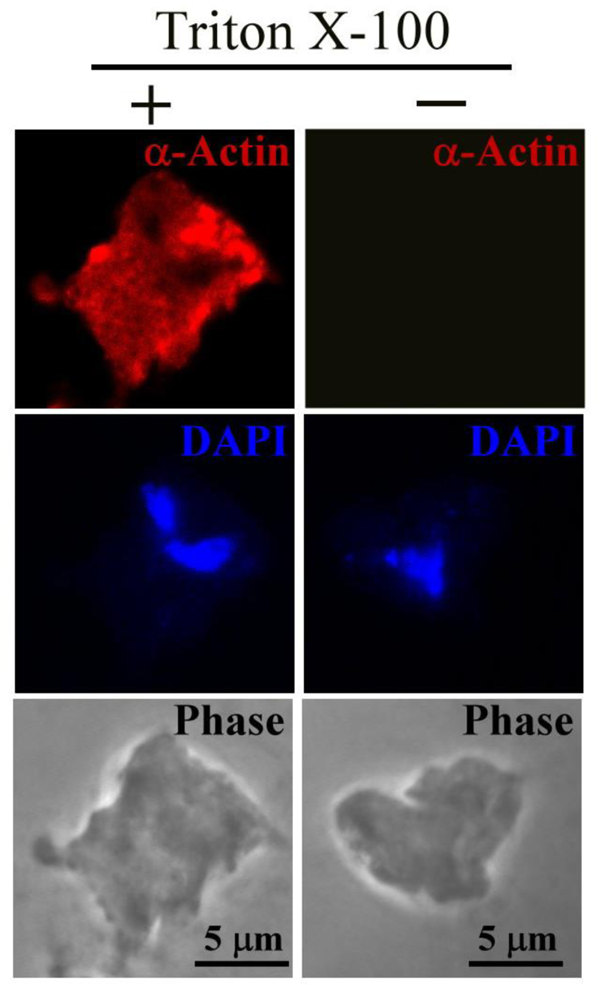
No detectable α-actin on the cell surface of *T. vaginalis*. The TH17 trophozoites with or without Triton X-100 permeation were examined by IFA using an anti-α-actin antibody, followed by incubation with a Cy3-conjugated secondary antibody. The nuclei were stained by DAPI. The morphology was observed by phase-contrast microscopy. The scale bars represent 5 μm.

**Figure 2-Figure Supplement 3.**
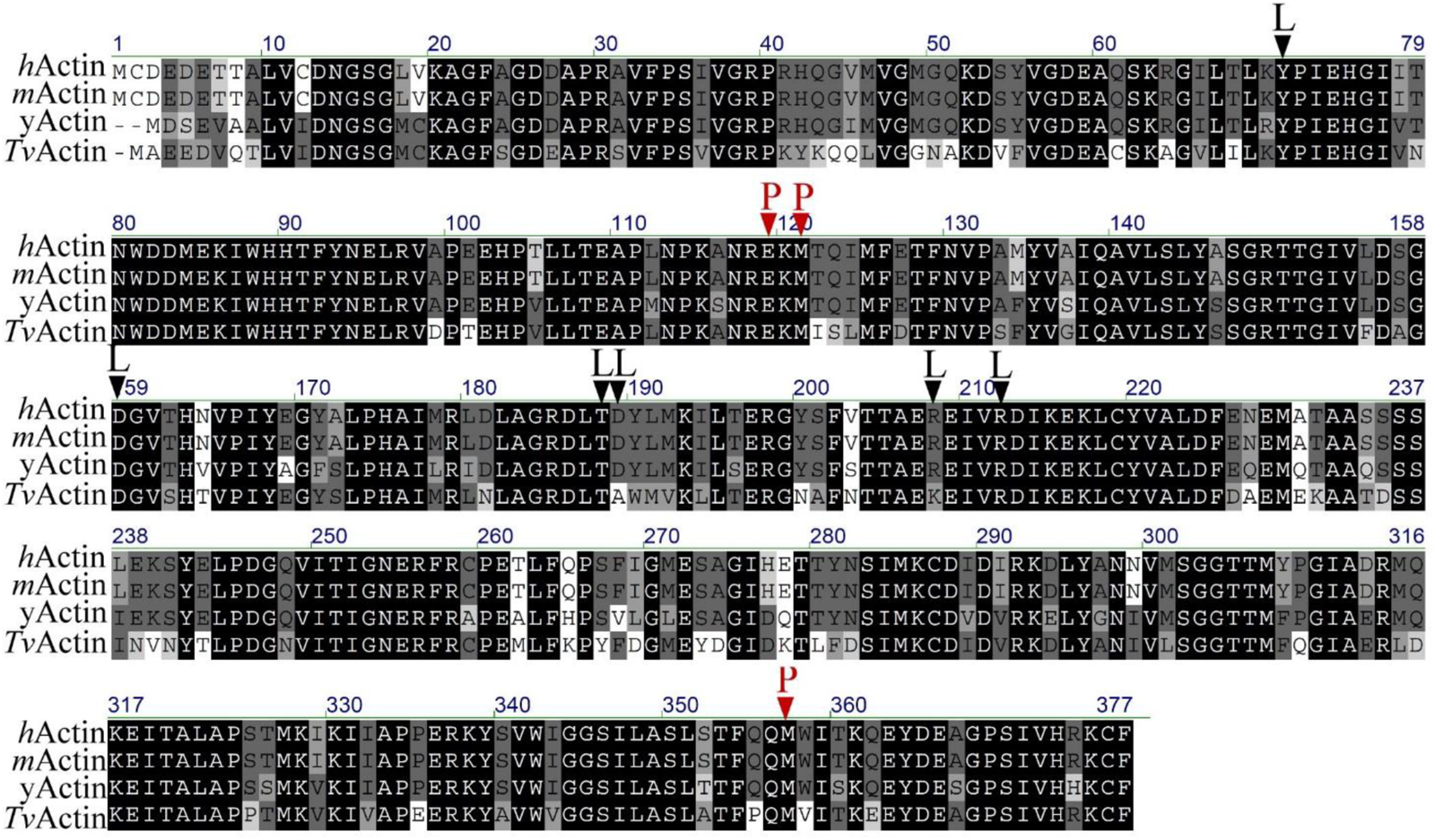
Protein sequence alignment of α-actin and *Tv*FACPα. (A.) The full-length protein sequences of α-actin from human (*h*Actin, P68133), mouse (*m*Actin, P68134), yeast (*y*Actin, P60010), and *T. vaginalis* (*Tv*Actin, TVAG_337240) were aligned to show the protein sequence similarity. The conserved amino acid residues are highlighted, and the binding sites of phalloidin or LatB in α-actin are indicated by P or L as shown at the top of sequences, respectively.

**Figure 4-Figure Supplement 1.**
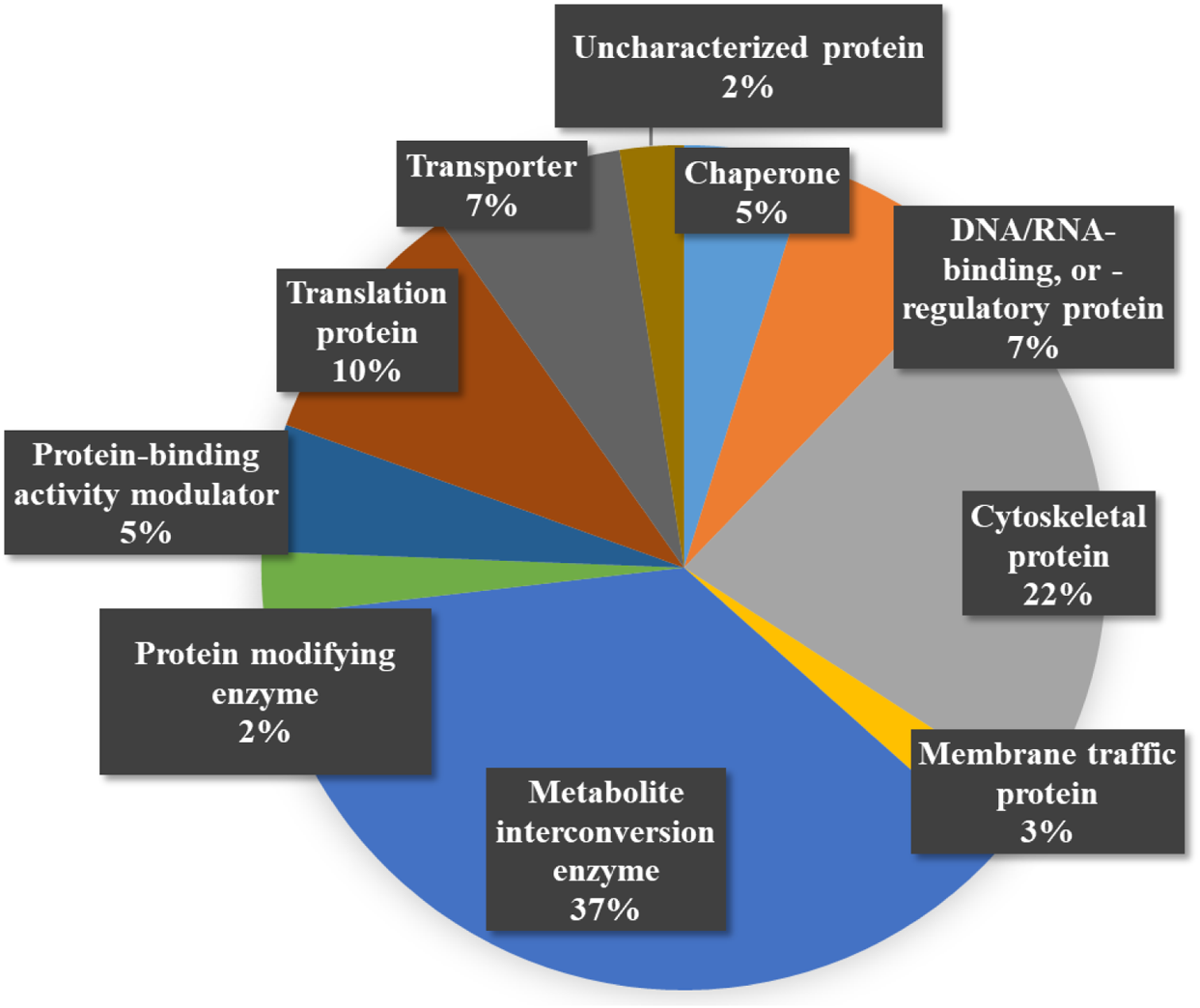
Protein function category. The mass identified proteins were classified by function into multiple cellular pathways, including cytoskeleton proteins (22%), chaperones (5%), membrane trafficking (3%), transporter (7%), protein binding (5%), modification (2%), DNA/RNA regulation (7%) and translation (10%), metabolism enzymes (37%), and uncharacterized proteins (2%).

**Figure 4-Figure Supplement 2.**
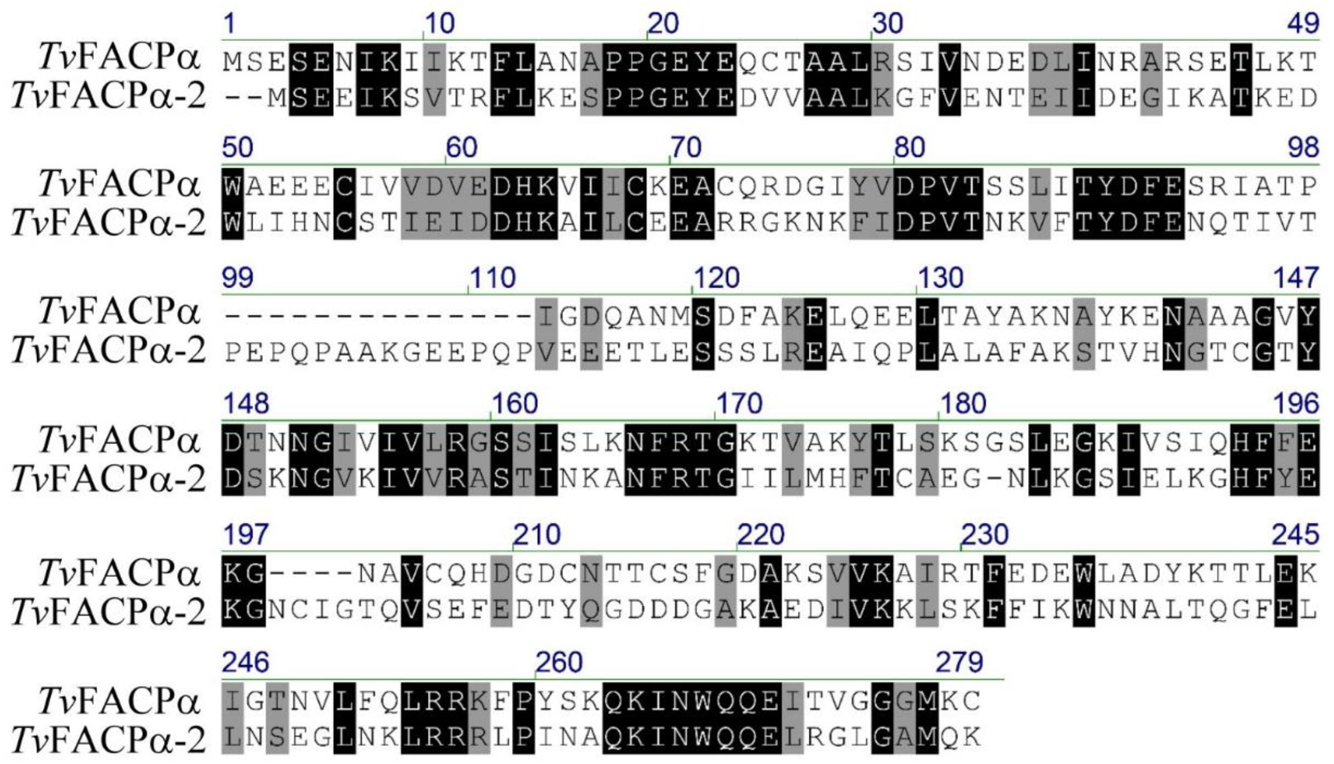
The sequence alignment for *Tv*FACPs in *T. vaginalis.* The protein sequences of *Tv*FACPα (TVAG_470230) and *Tv*FACPα-2 (TVAG_212270) were aligned. The conserved amino acid residues are highlighted.

**Figure 7-Figure Supplement 1.**
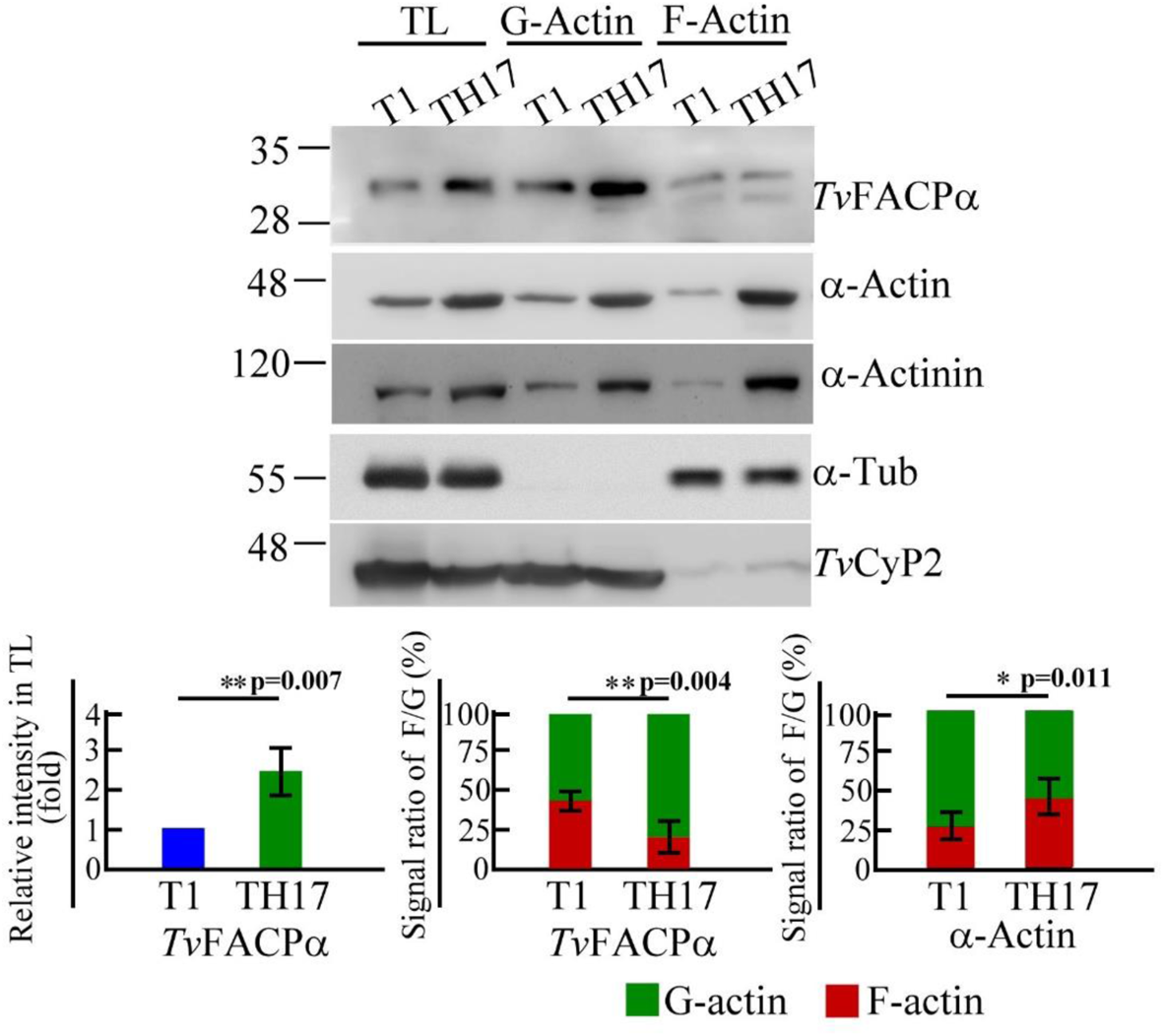
Differential expression of *Tv*FACPα in nonadherent and adherent isolates of *T. vaginalis.* The protein lysates from Figure 2E were re-examined by western blotting with the anti-*Tv*FACPα antibody. The relative intensity of *Tv*FACPα detected in total lysate, the signal ratio of *Tv*FACPα in F-actin versus G-actin fractions were shown in the histograms. The assays were repeated three times. Data in histograms are presented as mean ± SEM. Significance with p-value is statistically analyzed by Student’s t-test as indicated. (n=3, P<0.01, P<0.05, and ns, no significance).

**Figure 9-Figure Supplement 1.**
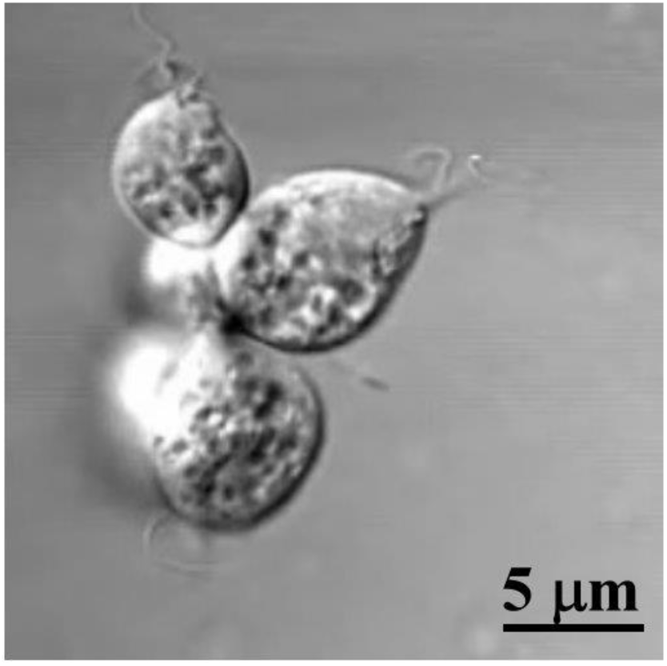
Morphology of trophozoites in the bottom well of trans-well assay. Morphology of trophozoites migrating into the bottom well was recorded by microscopy. The parasite in the bottom well were observed in dominant flagellate trophozoite with clear flagella under our assay conditions. The scale bar represents 5 μm.

## Notes

### Competing Interest Statement

The authors have declared no competing interest.

